# Immune Niche Formation in Engineered Mouse Models Reveals Mechanisms of Tumor Dormancy

**DOI:** 10.1101/2025.04.16.649000

**Authors:** Abdul Ahad, Feng Leng, Hiroshi Ichise, Edward Schrom, Jae Young So, Carter Sellner, Yang Gu, Wenjuan Wang, Celine Kieu, Woo Yong Park, Rachel Yang, Karen Wolcott, Ferenc Livak, Michael Kruhlak, Olga Aprelikova, Justin Gray, Vishal N. Kopardé, Yasuhiro Moriwaki, Ronald N. Germain, Li Yang

**Author notes:** Correspondence Building 37, Room 3134C 37 Convent Drive, Bethesda, MD 20892. **Financial support:** NCI intramural funding to Dr. Li Yang This work was supported in part by the DIR, NIAID, NIH [Lymphocyte Biology Section] and by a cooperative agreement between CCR, NCI and DIR, NIAID [Center for Advanced Tissue Imaging]. **Conflict of interest statement:** The authors declare no potential conflicts of interest.

## Abstract

Residual tumor cells can persist in a dormant state during clinical remissions that may last decades. The mechanisms that lead to such growth control vs. eventual reactivation and macroscopic tumor outgrowth remain unclear. Here, we report data from a mouse model that reveals a key role of host immunity and the cellular and molecular mechanisms that control tumor dormancy. Abrogation of myeloid-specific TGF-βRII expression (TβRII^myeKO^) resulted in an IFN-γ rich immune microenvironment. IFN-γ in turn elevated KLF4-mediated SLURP1 production in malignant cells, which is critical to the tumor cell quiescent state through interruption of fibronectin-integrin signaling pathways. The dormant tumor lesions were located in spatially localized immune niches rich in NK cells, cDCs, monocytes, and neutrophils, concomitant with tumor cell inactivation of NK cell immune surveillance through a CD200-CD200R1 mechanism. Our studies identify the IFN-γ-KLF4-SLURP1 and CD200-CD200R1 axes as critical molecular drivers in tumor dormancy regulated by immune-tumor crosstalk. These insights provide enhanced mechanistic understanding of tumor dormancy in a mouse model suitable for further investigation of cancer treatment resistance and prevention of metastatic spread.

## Introduction

Residual tumor cells can persist in a dormant state during extended periods of clinical remission ^1–5^, with the ‘reawakening’ of dormant tumor cells after months to years often resulting in incurable metastatic outgrowth and patient mortality ^1, 2, 6–8^. Such quiescence has been recognized as a key mechanism by which tumor cells can evade host immune detection, clearance, and therapeutic targeting ^5, 9–13^ but progress in understanding the mechanisms governing tumor dormancy has been limited ^5, 8, 14^.

Tumor dormancy can be characterized as cellular dormancy, that is, singular or a small cluster of quiescent cells without ongoing cell proliferation ^15, 16^, or tumor mass dormancy, which involves micro-metastatic nodules that maintain an equilibrium between cell proliferation and intrinsic or immune-mediated death with no net increase in size over an extended period of time ^1, 8, 17–21^. The quiescent state in cellular dormancy offers not only protection from many conventional chemotherapies that rely on cell cycle progression to exert their cytotoxic effect but also promotes evasion of the host immune system ^12, 22^. Several mechanisms regulating cellular dormancy have been identified, including those involving signaling pathways (e.g., TGF-β2, ERK1/2, and P-p38) ^23, 24^, nuclear receptor NR2F1 ^25–27^, stemness factors ^10, 28, 29^, and extracellular matrix mediators ^30–35^. In addition, recent studies support the ability of dormant tumor cells to evade CD8^+^ T cell– and natural killer cell (NK cell)–mediated detection and clearance ^12, 21, 29, 36–39^, with constraint of reactivated metastatic cells being analogous to control of latent / chronic viral infection with cytomegalovirus or herpes simplex virus. With these pathogens, episodic activation of latent virus is immediately targeted by the adaptive immune system to prevent overt viremia ^40–42^ – in the case of cancer, such constant immunosurveillance and anti-tumor effector responses may prevent macroscopic tumor formation over an extended period, with eventual immune escape accounting for development of clinically relevant metastatic disease ^19, 43^.

At present there is a lack of mechanistic insight into how disseminated tumor cells enter a reversible growth-arrest or quiescence state and what casual roles immune cells and molecular mediators play in keeping dormant cancer cells from being reactivated or developing into a clinically relevant tumor ^44^. Additionally, it is unclear whether distinct immune niches, either microenvironments that are pre-organized by the host or ones created by disseminated tumor cells, actively facilitate tumor dormancy ^4, 8^. One of the major challenges in filling these knowledge gaps is the lack of immune-competent mouse models that can effectively produce a long-term tumor dormancy phenotype *in vivo*, especially in the context of immune regulation ^44 5, 45^.

Our studies over the last two decades have uncovered a key role for myeloid TGF-β signaling in promoting tumor immune evasion and metastatic outgrowth ^46–49^. Here we use this model with a combination of advanced imaging, transcriptomics, and molecular studies to gain deeper insights into how modulation of myeloid cell innate immune state can prevent tumor outgrowth. We show that abrogation of myeloid TGF-β signaling induced an IFN-γ rich immune milieu, leading to KLF4-SLURP1 upregulation that facilitates the tumor dormant state. The microdomains of dormant tumors contained accumulations of myeloid and lymphoid immune cells and this microdomain modulation is complemented by resistance of the tumor cells to NK cell-mediated killing via CD200-CD200R1 interactions. Together, these results show how the immune system can promote prolonged maintenance of viable metastatic tumor cells by facilitating entry into the dormant state in concert with immune escape mechanisms. Consistent with these findings, human correlative studies reveal a poor prognosis and early post-treatment relapse associated with tumor cell IFN-γ-KLF4-SLURP1 and CD200-CD200R1 gene signatures. Our results highlight the Tgfbr2^MyeKO^ mouse as a valuable tool for studying tumor dormancy, opening avenues for further investigation into dormant cancer cell immune evasion and strategies to prevent metastatic outgrowth.

## Results

### Abrogation of myeloid specific TGF-β signaling induces cellular tumor dormancy in multiple mouse models of breast cancer metastasis

We have previously reported that deletion of *Tgfbr2*, the gene encoding TGF-β receptor II (TβRII) in myeloid cells (Tgfbr2^MyeKO^) decreases macroscopic cancer metastasis through increased host anti-tumor immunity ^47, 49^. In the current study we employed D2A1, a proliferative isogenic breast cancer cell line that is widely used for tumor dormancy studies ^15, 50^ to explore whether such resistance to macroscopic tumor growth involves clearance of cancer cells or induction of dormancy, with the potential for future outgrowth. D2A1 cells were transduced with the Doxycycline (Dox) inducible H2B-GFP system, allowing them to be used in immune-competent mice and for microscopic imaging. Using this model, we find a striking cellular dormancy phenotype upon abrogation of myeloid-specific TGF-β signaling. The dormant tumor lesions are characterized by single cells or cell clusters with fewer than 8 cells (1-8 cell per lesion) in Tgfbr2^MyeKO^ mice 12, 30, and 50 days after tail vein injection (TVI) of D2A1 tumor cells (Figure 1A). This is approximately the equivalent of 1.5-5.5 years in humans, a relevant time window in the context of triple-negative breast cancer relapse (TNBC) and immune checkpoint blockade (ICB) resistance. Consistent with our previous publications, the Tgfbr2^MyeKO^ mice showed fewer gross metastatic nodules and increased survival after TVI with D2A1 cells (Supple fig. S1A-B). The dormancy phenotype was also observed in the TSAE1+mHer2 breast cancer mouse model (Figure 1B) as well as in the 4T1 orthotopic model, an aggressive breast cancer metastatic model where increased numbers of dormant lesions and decreased proliferative lesions were evident in the Tgfbr2^MyeKO^ mice (Figure 1C).

**Figure 1.**
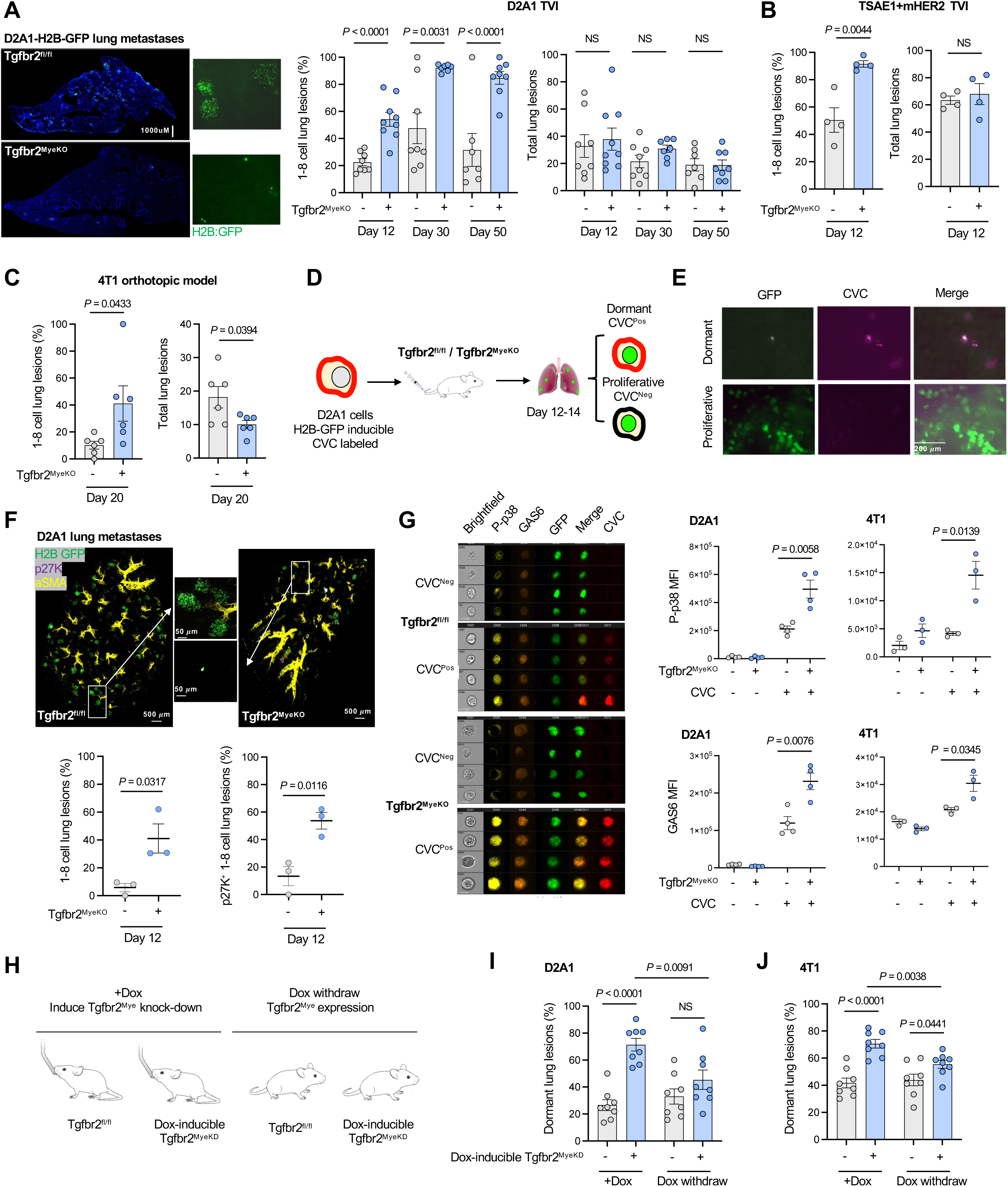
A mouse model of tumor dormancy induction upon abrogation of myeloid-specific TGF-β signaling. **A.** Representative lung surface imaging of metastatic lesions from Tgfbr2^MyeKO^ and Tgfbr2^fl/fl^ (flox cont.) mice that received D2A1-H2B-GFP TVI for 12-14 days (left), and % dormant lesions (<8 cells) (middle) and total tumor lesions (right) at 12, 30, and 50 days after mice received TVI of tumor cells. n=7-9 mice per group. **B.** % 1-8 cell tumor lesions (left) and total tumor lesions (right) from the TSAE1+mHer2 TVI model, imaging on day 12. n=4 mice per group. **C.** % 1-8 cell tumor lesions (left) and total tumor lesions (right) from 4T1 orthotopic model, imaging on day 20. n=6 mice per group. **D.** Schematic design for sorting dormant (GFP+CVC^Pos^) and proliferative D2A1-H2B-GFP (GFP+CVC^Neg^) cells. **E.** Representative lung surface images of dormant (GFP+CVC^Pos^) and proliferative lesions (GFP+CVC^Neg^) from mice that received TVI of D2A1-H2B-GFP for 12 days. **F.** Representative Clearing-enhanced 3D (Ce3D) confocal imaging of lung tissues at 12 days after TVI of D2A1-H2B-GFP-mRuby-p27K cells. Green: H2B-GFP, Violet: mRuby-p27, and yellow: aSMA, with enlarged images in the middle panels; % 1-8 cell dormant lesions (lower left) and p27K+ dormant lesions (lower right). n=3 mice per group. **G.** Representative Imaging flow cytometry of P-p38 and GAS6 in dormant GFP+CVC^Pos^ and proliferative GFP+CVC^Neg^ D2A1 cells from Tgfbr2^MyeKO^ and flox cont. mice (left) and average of fluorescence intensity of P-p38 and GAS6 in GFP+CVC^Pos^ and GFP+CVC^Neg^ D2A1 and 4T1 cells (right). n=3-4 mice per group. **H**. Schematic design for the effect of Dox-induced myeloid-TβRII knockdown (KD) and myeloid-TβRII re-expression on dormant lesions. **I-J.** % dormant lesions from TβRII KD and re-expression after D2A1 TVI (I) and 4T1 orthotopic injection at day 12 (J). n=8 mice per group. All data are presented as mean ± s.e.m. *P*-values were derived from a two-tailed Student’s t-test and significance was determined by a *P*-value < 0.05.

To further characterize dormant lesions from the Tgfbr2^MyeKO^ mice, we labeled D2A1 cells with Cellvue Claret (CVC), a lipophilic dye that is embedded in the plasma membrane and is not degraded during protein turnover but diminishes in intensity upon cell division (Figure 1D). Solitary dormant D2A1 cells retained CVC labeling, while micro-metastatic lesions did not (Figure 1E). Additionally, we transduced D2A1 cells with the mRuby-p27K quiescence reporter which lacks Cyclin-dependent kinase (CDK) inhibitory activity but retains the susceptibility for ubiquitin-mediated degradation, enabling visualization of cells in the G0 phase. Clearing-enhanced 3D (Ce3D) volume confocal imaging of the lung tissue revealed that approximately 60% of 1-8 cell lung lesions in Tgfbr2^MyeKO^ mice were p27K positive (Figure 1F). Further, imaging flow cytometry analysis revealed higher expression of the previously reported tumor dormancy markers phosphorylated p38 (P-p38) and GAS6, or the two in combination ^44, 51, 52^, in CVC^Pos^ dormant cells from Tgfbr2^MyeKO^ mice as compared to those from controls (Figure 1G, and Supple fig. S1D-E). Based on these data, the 1-8 cell clusters will be referred to as dormant tumor lesions.

We next investigated whether rescue of myeloid TβRII expression would diminish the dormancy phenotype using a Dox inducible myeloid TβRII shRNA knockdown transgenic mouse line (Figure 1H). As expected, Dox-induced myeloid TβRII knockdown before D2A1 TVI or 4T1 orthotopic implantation increased the percentage of dormant lesions while withdrawing Dox decreased the frequency of such lesions (Figure 1I, 1J). In contrast, Dox-induced myeloid TβRII knockdown after D2A1 TVI did not increase the frequency of dormant lesions over control (Supple fig. S1G), suggesting the importance of a pre-established distant organ microenvironment in the formation of dormant lesions. There were no changes in the total number of tumors (Supple fig. S1F and G), indicating that cell death of founder metastatic cells in the lung microenvironment of the Tgfbr2^MyeKO^ mice did not play a major role in these findings. Together, these data further support the value of Tgfbr2^MyeKO^ mice as a relevant *in vivo* model of tumor dormancy using multiple mouse breast cancer cell lines and reveal that while suppressing gross tumor outgrowth, the loss of TGFβ signaling in myeloid cells promotes survival of metastatic tumor cells in a dormant state.

### SLURP1-mediated induction of the tumor cell quiescent state

To gain insight into the molecular features of the dormancy state upon abrogation of myeloid TGF-β signaling, we sorted GFP+CVC^Pos^ dormant and GFP+CVC^Neg^ proliferating tumor cells from control and Tgfbr2^MyeKO^ mice 14 days after TVI of D2A1-H2B-GFP cells (Figure 2A, and Supple fig. S2A, S2B, S2C, S2D). RNA-seq analysis (biological triplicates: n=50 each for dormant and proliferative cells) showed four distinct cell clusters in the PCA plot (Figure 2A). By comparing dormant and proliferating D2A1 cells from control and Tgfbr2^MyeKO^ mice, we identified 504 differentially expressed genes that were unique to dormant cells from Tgfbr2^MyeKO^ mice (Figure 2B). As expected, gene set enrichment analysis (GSEA) of dormant cancer cells revealed a decreased enrichment of cell cycle mediators including targets of E2F and MYC, as well as mTOR signaling (Figure 2C).

**Figure 2.**
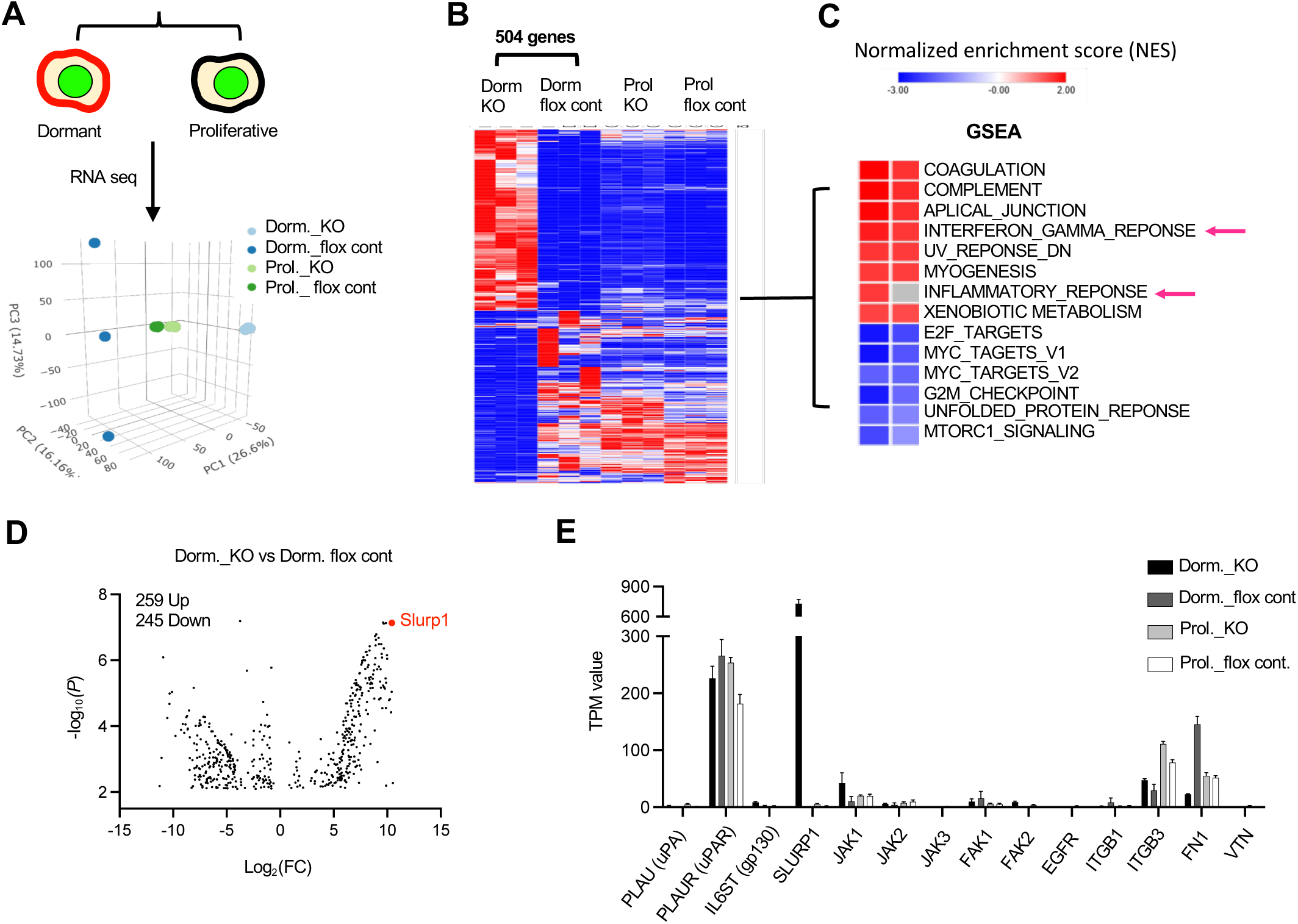
Increased SLURP1 expression in GFP+CVC^Pos^ dormant cells from the Tgfbr2^MyeKO^ mice. **A.** Schematic for sorting dormant and proliferative tumor cells using inducible H2B-GFP and CVC membrane labeling (left) and PCA plot from RNA-seq analysis (right). Dorm: sorted dormant tumor cells; Prol: sorted proliferative tumor cells; flox cont. mice; KO: Tgfbr2^MyeKO^ mice. n=3 mice per group. **B.** Heatmap of 504 differential genes from RNAseq data analysis comparing dormant vs proliferative tumor cells from Tgfbr2^MyeKO^ and flox cont. mice. n=3 mice per group. **C.** Gene set enrichment analysis (GSEA) of differential genes shown in B. **D.** Volcano plot for upregulated and downregulated genes comparing dormant cells from the Tgfbr2^MyeKO^ with those from flox cont. mice, with SLURP1 (in red) among the top upregulated. **E.** TPM values of genes from RNA-seq revealing increased SLURP1 and integrin-mediated pathway including Itgb1, Fn1, uPA and uPAR. n=3 mice per group.

Slurp1 was one of the most upregulated genes in dormant cells from Tgfbr2^MyeKO^ mice (Figure 2D and 2E). SLURP1, a secreted Ly6/uPAR related protein, disrupts uPA (urokinase plasminogen activator)-mediated cell proliferation in corneal homeostasis ^53, 54^ and shows anti-proliferation properties in cancer ^55^. Imaging flow cytometry revealed a higher expression of SLURP1 in CVC^Pos^ D2A1 and 4T1 dormant cells from Tgfbr2^MyeKO^ mice as compared to tumor cells from control mice (Figure 3A). These SLURP1^+^CVC^Pos^ cells also showed high levels of P-p38 and GAS6 (Figure 3B and Supple fig. S3A), consistent with our findings above.

**Figure 3.**
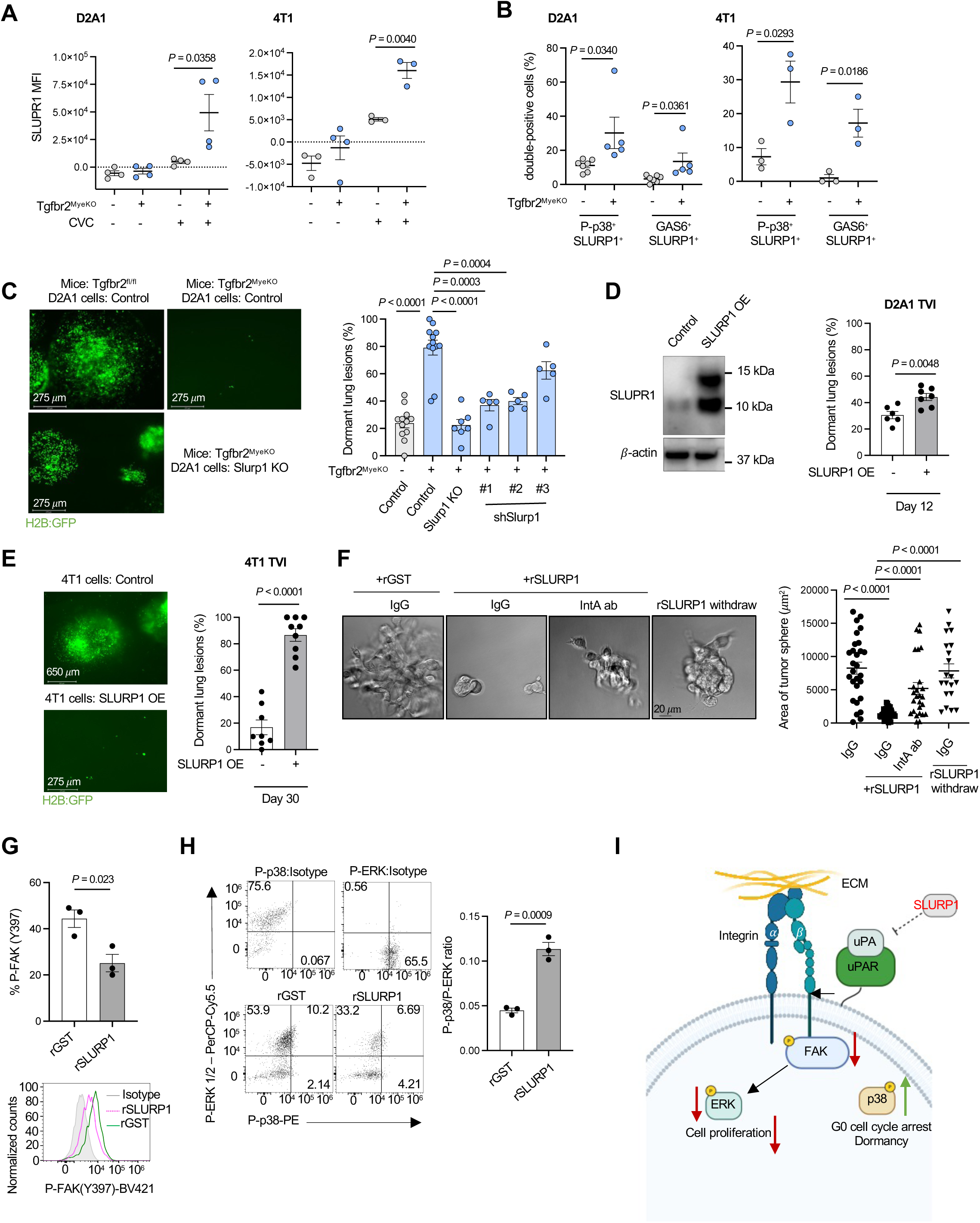
The effect of SLURP1 on tumor dormancy. **A.** SLURP1 fluorescence intensity in dormant GFP+CVC^Pos^ and proliferative GFP+CVC^Neg^ tumor cells from Tgfbr2^MyeKO^ and flox cont. mice by Imaging flow cytometry, D2A1 (left) and 4T1 (right); n=4 biological independent experiments. **B.** % of SLURP1^+^ cells with P-p38^+^ and GAS6^+^ expression from dormant GFP+CVC^Pos^ and proliferative GFP+CVC^Neg^ tumor cells from Tgfbr2^MyeKO^ and flox cont. mice; n=4 four biological independent experiments. **C.** Representative lung surface imaging (left), and % dormant lesions from D2A1 tumor cells with Slurp1 KO and KD (right). n=5-12 mice per group. **D.** Western blot for SLURP1 overexpression (SLURP1-OE) in D2A1 tumor cells (D2A1-SLURP1-OE) (left), and % dormant lesions from mouse lungs at 12 days after TVI of D2A1-SLURP1-OE tumor cells (right). **E.** % dormant lesions from mouse lungs at day 30 after TVI of 4T1-SLURP1-OE tumor cells. n=9-10 mice per group. **F.** Representative images of D2A1 spheroid culture treated with recombinant SLURP1 (rSLURP1), and rSLURP1 plus an integrin activating antibody (IntA ab), as well as with rSLURP1 withdraw (left) and quantification of spheroid size (right). **G-H.** Flow cytometry analysis of P-FAK (G), P-p38, P-ERK, and the ratio of P-p38 to P-ERK (H) in 3D-cultured D2A1 cells treated with rSLURP1 or GST control. **H.** Schematic for SLURP1 mediated inhibition of tumor cell proliferation (created with BioRender.com). All data are presented as mean ± s.e.m. *P*-values were derived from a two-tailed student t-test. Significance was determined by a *P*-value< 0.05.

To investigate the contribution of SLURP1 to the dormant state we generated Slurp1 knock-out (KO) and Slurp1 shRNA knockdown (KD) D2A1 cells (Supple fig. S3B, S3C). Tgfbr2^MyeKO^ mice injected with Slurp1 KO or KD tumor cells showed fewer dormant lesions (Figure 3C) with only a modest effect on the total tumor lesion number (Supple fig. S3D). Conversely, SLURP1 overexpression in D2A1 and 4T1 cell lines increased the percentage of dormant lesions in wild-type mice (Figure 3D, 3E and Supple fig. 3E), accompanied by decreased total lesions (D2A1) or no difference in lesion number (4T1) (Supple fig. S3F). Consistently, D2A1 with SLURP1 overexpression decreased the number of gross metastatic nodules quantified by Indian Ink staining (Supple fig. S3G). In *in vitro* 3D culture, recombinant SLURP1 treatment decreased D2A1 spheroids size in a dose-dependent manner (Supple fig. S3H).

SLURP1 has been reported to interact with uPA ^53^ and uPA-uPAR interaction is critical to the activation of Integrin-FAK-ERK signaling pathways and promotion of tumor cell proliferation ^52, 56, 57^. When an integrin-activating antibody (IntA ab) was added to the D2A1 spheroid culture, it reversed the inhibitory effect of recombinant SLURP1 on spheroid growth (Figure 3F), and as expected, withdrawing SLURP1 treatment rescued spheroid growth (Figure 3F). This connection of SLURP1 expression to adhesion and tumor growth was also supported by RNA-seq analysis of tumor spheroids treated with recombinant SLURP1 compared with GST control, which showed integrin cell surface interactions and extracellular matrix organization as the top enriched pathways (Supple fig. S3I). Further, recombinant SLURP1 treatment decreased FAK phosphorylation (Figure 3G) and increased the P-p38 to P-ERK ratio (Figure 3H), characteristic of dormant cancer cells ^58^. Together with previous findings in the literature, our *in vivo* and *in vitro* data support an inhibitory role of SLURP1 in tumor cell proliferation by targeting the Integrin-FAK-ERK signaling pathways (Figure 3I).

### IFNγ-KLF4 axis regulation of SLURP1 expression

What features of the tumor microenvironment in Tgfbr2^MyeKO^ mice promoted the increased expression of SLURP1 that limits tumor proliferation? Examination of differentially-expressed genes comparing Tgfbr2^MyeKO^ dormant tumor cells vs. those from the control mice suggested that dormancy may arise from signals within an activated immune microenvironment dependent on myeloid-specific *Tgfbr2* deletion (Figure 2C, red arrows). Further, a Bio-Plex immunoassay detected higher levels of IFN-γ as well as TNF-α in the lung microenvironment of tumor-bearing Tgfbr2^MyeKO^ mice as compared to control mice (Supple fig. S4A). Recombinant IFN-γ treatment of D2A1 tumor cells increased SLURP1 mRNA *in vitro* under stress and nutrient-deprived conditions (Figure 4A). Importantly, KD of the IFN-γ receptor (IFN-γR) decreased SLURP1 expression in D2A1 cells (Figure 4B and C, Supple Fig. S4B), and this was associated with decreased P-p38 but not GAS6 protein expression in CVC^Pos^ D2A1 cells from Tgfbr2^MyeKO^ mice (Supple Fig. 4C). IFN-γR KD in D2A1 cells also decreased the percentage of dormant lesions in Tgfbr2^MyeKO^ mice (Figure 4D). These data suggest the importance of IFN-γ signaling in promoting SLURP1 expression and tumor dormancy induction.

**Figure 4.**
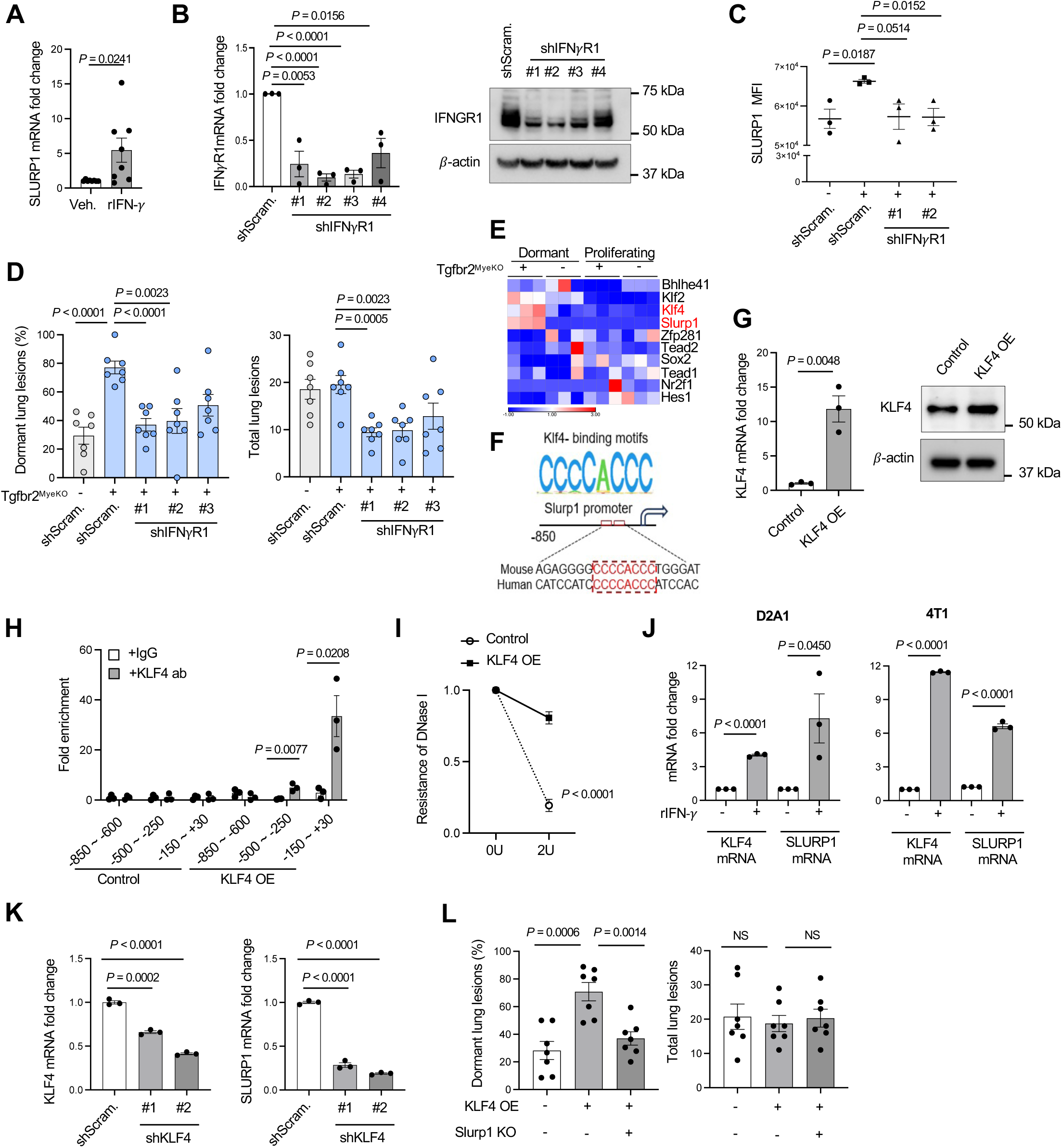
IFN-γ-KLF4 axis regulation of SLURP1 expression. **A.** SLURP1 expression in cultured tumor cells treated with IFN-γ. n=8, independent biological replicates. **B.** RT-qPCR and Western blot validation of IFN-γR KD in D2A1 cells. **C.** SLURP1 expression from Imaging flow cytometry of dormant cells with or without IFN-γR KD, single cell suspension from the lungs of the tumor-bearing mice. n=3 biologically independent experiments. **D.** Dormant (left) and total tumor lesions (right) from lung surface imaging of mice that received D2A1 cells with or without IFN-γR KD. n=7-8 mice per group. **E.** TPM expression heatmap of KLF4 transcription family members from RNA-seq of the dormant cells. n=3 mice per group. **F.** KLF4 binding site mapping in the *Slurp1* promoter. Putative KLF4-binding motifs are predicted in the mouse *Slurp1* promoter(up) by Homer assay. Klf4 binding motif CCCCACCC were shown in mouse and human *Slurp1* promoter. **G.** RT-qPCR and Western blot of KLF4 overexpression in D2A1 cells. **H.** Fold enrichment of SLURP1 expression from KLF4-ChIP qPCR of D2A1 cells, n=3 biologically independent experiments. Results shown as mean ± SD. **I.** DNase-mediated DNA degradation showing KLF4 overexpression protected the KLF4 binding regions in *Slurp1* promoter. n=3 biologically independent experiments. Results are shown as mean ± SD. **J.** RT-qPCR of KLF4 and SLURP1 expression in cultured tumor cells treated with recombinant IFN-γ (left: 4T1, and right: D2A1). **K.** The effect of KLF4 KD (left) on fold changes of SLURP1 expression (right) in D2A1 cells. n = 3 experiments. **L.** Dormant (left) and total tumor lesions (right) from lung surface imaging of mice that received TVI of D2A1 tumor cells with KLF4 OE, as well as D2A1 cells with KLF4 OE plus Slurp1 KO. n=7 for each group. Data are presented as mean ± s.e.m. *P*-values were derived from a two-tailed Student’s t-test. Significance was determined by a *P*-value< 0.05.

Using data from our RNA-seq analysis comparing dormant and proliferating cells, we next looked for differential expression of transcription factors that might regulate SLURP1 expression and found KLF4 and KLF2 but not other Krüppel-like factor family members were increased in D2A1 cells from Tgfbr2^MyeKO^ mice (Figure 4E). The Homer assay predicted a putative KLF4-binding motif CCCCACCC to be present in mouse and human Slurp1 promoters (Figure 4F). KLF4-ChiP assays in D2A1 tumor cells with KLF4 overexpression (Figure 4G) revealed that KLF4 protein was enriched at the -500∼-250 and −150∼+30 bp regions of the SLURP1 promoter (Figure 4H), consistent with this motif mapping. Moreover, the KLF4 DNA binding sites in the SLURP1 promoter were protected from DNA degradation in D2A1 dormant cells (Figure 4I). In accord with these findings, both KLF4 and SLURP1 were increased in 4T1 and D2A1 tumor cells upon treatment with IFN-γ (Figure 4J). Additionally, shRNA KD of KLF4 decreased SLURP1 expression and KLF4 overexpression increased it in D2A1 cells (Figure 4K and Supple fig. S4D). In wild-type mice, the D2A1 cells with KLF4 overexpression showed increased numbers of dormant lesions and Slurp1 KO in these KLF4 overexpression cells reversed the dormancy phenotype (Figure 4L), indicating SLURP1 indeed serves as a downstream mediator of KLF4’s dormancy effect. Together, these results suggest a critical role of an IFN-γ-KLF4-SLURP1 axis in the observed tumor dormancy phenotype.

### Elevated IFN-γ levels in metastatic lung tissue and immune niches of the Tgfbr2^MyeKO^ mice

Myeloid *Tgfbr2* deletion alters immune cell function, with a significant impact in both tumor-bearing mice and stroke occurrence in aging mice ^47, 59^. To better understand the way in which the *Tgfbr2* deletion affected local immune events that contributed to the observed tumor dormancy phenotype, we used spectral flow cytometric profiling of immune cells in the lungs of tumor-bearing Tgfbr2^MyeKO^ and control mice. The primary change among myeloid cells was an increase in CD103^+^ dendritic cells (DCs) (Figure 5A and 5B), with no difference in the overall number of CD45^+^ immune cells (Supple fig. S5A, S5B). In addition, the expression of TNF-α was significantly higher in CD103^+^ DCs from tumor-bearing Tgfbr2^MyeKO^ mice (Figure 5C). In the lymphoid compartment, tumor-bearing Tgfbr2^MyeKO^ mice showed a substantial increase in IFN-γ^+^CD4^+^ and especially IFN-γ^+^CD8^+^ T cells, and a more modest but significant increase in IFN-γ^+^ NK cells (Figure 5D, Supple fig. S5C). These data suggest an increase in type 1 (inflammatory) immunity at the metastatic lung site in the Tgfbr2^MyeKO^ mice.

**Figure 5.**
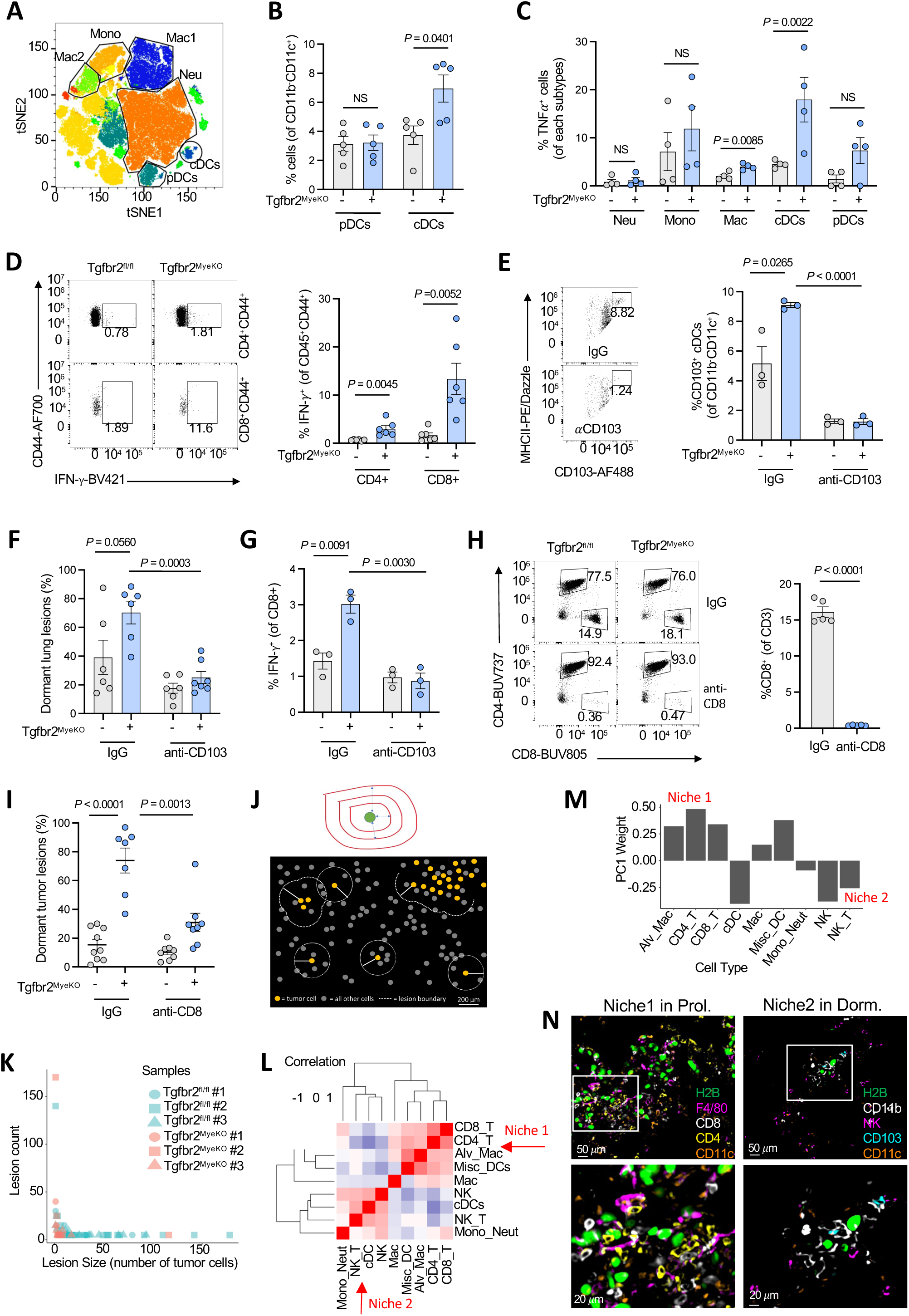
Elevated IFN-γ levels in metastatic lung tissue and immune niches of the Tgfbr2^MyeKO^ mice. **A.** tSNE plots for all myeloid cells by Cytek analysis. **B.** % CD103^+^ DCs and plasmacytoid DCs (pDCs) in CD45^+^CD11b^-^ population. **C.** %TNF-α^+^ neutrophiles, monocytes, macrophages, CD103^+^ DCs and pDCs. n=5-6 mice per group. **D.** Flow cytometry analysis of IFN-γ^+^CD4 and IFN-γ^+^CD8 T cells (gated on CD44^+^ effector cells) from the lung of Tgfbr2^MyeKO^ and flox cont. mice. n=7 mice per group. **E.** Flow cytometry validation of CD103^+^ DC depletion. **F and G.** Tumor lesions in the lungs (F) and Cytek analysis of IFN-γ^+^CD8^+^ T cells (G) from the lungs of Tgfbr2^MyeKO^ and flox cont. mice upon CD103^+^ DC depletion. n=3 mice per group. **H.** Flow cytometry validation of CD8 T cell depletion from the lung of Tgfbr2^MyeKO^ and flox cont. mice. **I.** Tumor lesions in the lungs of mice with CD8 T cell depletion. n =7 mice per group. **J.** Strategy for demarcating individual tumor lesions separated by 200 µm distance to define neighbors. n=3 each for Tgfbr2^MyeKO^ and flox cont. mice. **K.** Distribution of tumor lesion sizes from each mouse lung sample. flox cont. lesions are larger on average (zero-truncated negative binomial regression model, *P* = 0.0028). **L.** Pairwise correlations of standardized center log ratios (CLRs) for each immune cell type across all tumor lesions. Two immune niches of commonly co-occurring cell types are outlined and indicated. **M.** Loadings for PC1 of the PCA, where a positive weight indicates increasing fractional abundance of a cell type as PC1 increases, and a negative weight indicates decreasing fractional abundance of a cell type as PC1 increases. **N.** Representing images showing niche 2 (CD103^+^cDCs, NK cells Monocyte/Neutrophils) surrounding dormant tumor lesions in Tgfbr2^MyeKO^ mouse lungs and niche 1 (Alv macrophages, CD8^+^ and CD4^+^ T cells) surrounding proliferative tumor lesions. All data are presented as mean ± s.e.m. *P*-values were derived from a two-tailed student t-test. Significance was determined by a *P*-value< 0.05.

To directly interrogate the role of cDC1 and CD8 T cells in driving the dormancy phenotype in Tgfbr2^MyeKO^ mice, we first depleted CD103^+^ DCs. Depletion of CD103^+^ DCs but not CD317^+^ pDCs decreased dormant lesions (Figure 5E and F, and Supple fig. S5D-5F), suggesting a likely contribution of cDC1 to the observed tumor dormancy phenotype. Additionally, depletion of CD103^+^ DCs decreased IFN-γ producing CD8^+^ T cells in Tgfbr2^MyeKO^ mice (Figure 5G), with no effect on IFN-γ^+^CD4 T cells or TNF-α^+^ CD4 or TNF-α^+^CD8 T cells (Supple fig. S5G). Furthermore, as expected and consistent with our previous findings ^47^, CD8^+^ T cell depletion increased proliferative lesions and concurrently diminished the Tgfbr2^MyeKO^ tumor dormancy phenotype (Figure 5H and I, and Supple fig. S5H). IFN-γ was the critical mediator in this dormant tumor phenotype as neutralizing IFN-γ antibodies, but not antibodies to TNF-α, decreased CVC^Pos^ and increased Ki-67^+^ D2A1 cells in a coculture of CD103^+^DCs and CD8^+^ T cells from Tgfbr2^MyeKO^ mice with CVC labeled D2A1 tumor cells (Supple fig. S5I).

cDC1 and CD8^+^ T cells are primary components in the TME of effective adaptive anti-tumor responses, especially upon ICB immunotherapy; their possible involvement in promoting development of immune-resistant dormant tumor cells responsible for late arising metastatic lesions in patients with apparent remissions is thus a critical issue to examine, as it could reveal a duality to the effects of immunotherapy. We therefore next asked whether dormant tumor lesions resided in a distinct cellular niche where they escaped immune clearance as compared to sites of tumor cell proliferation. We employed IBEX multiplex immunostaining/imaging technology ^60^ (Table 3) and a custom pipeline for defining tumor lesions and their immune niches (Figure 5J). We segmented and classified all cells in the IBEX images based on known markers, defined lesions as groups of tumor cells separated by no more than 200μm and included all immune cells within 200μm of any tumor cell as part of that lesion (Figure 5J and Supple fig. S6A-B). We identified 183 tumor lesions, 115 from control and 68 from Tgfbr2^MyeKO^ mice. As expected and consistent with the dormancy phenotype upon abrogation of myeloid TGF-β signaling, tumor lesions were significantly larger in control mice (*P* = 1.5*10^-10^), containing an average of 16.8 tumor cells compared to an average of 3.8 tumor cells in the lesions of Tgfbr2^MyeKO^ mice (Figure 5K). Next, we used pairwise abundance correlations (Figure 5L) and principal component analysis (PCA) (Figure 5M and Supp Figure S6C) to study the immune cell compositions of the tumor lesions. These approaches independently uncovered two distinct immune niches: Niche 1 contains CD4^+^ and CD8^+^ T cells, alveolar macrophages, macrophages, and diverse DCs; surprisingly, these sites containing the T cells shown above to be the major producers of IFN-γ were associated with small proliferative lesions rather than cellular dormancy; conversely, Niche 2 was enriched in CD103^+^ cDCs, NK cells, NKT cells, and monocytes/neutrophils (Supp Figure S6D and S6E) and associated with true tumor cell dormancy. Smaller tumor lesions tended to have Niche 2-like immune cell compositions (*P* = 0.0006), as did lesions from Tgfbr2^MyeKO^ mice (*P* = 0.0190), while larger tumor lesions and tumor lesions from control mice tended to have Niche 1-like immune cell compositions (Supp Figures S6D and S6E, shown visually in Figure 5N). These spatial findings suggest a set of events are involved in the generation of the IFN-γ that induces dormancy, which protects tumor cells from immune attack, and constrains tumor lesions from proliferative expansion. They suggest a model in which a subset of metastatic tumor cells able to be rapidly induced to dormancy by IFN-γ upon seeding to the lung (possibly produced at a distance by activated CD8 T cells) escape destruction by nearby NK cells, whereas those tumor cells that avoid induction of dormancy by IFN-γ proliferate but are kept as small proliferating lesions by persistent cytotoxic action of the colocalized CD8 T cells.

### CD200-CD200R1 mediated immune evasion of dormant tumor cells

To better dissect this complex interplay between immune cells and tumor state, we focused on possible evasion of killing of dormant cells in Niche 2 containing many NK cells. Our RNA-seq data revealed several potential immune mediators and regulatory pathway candidates (Figure 6A, Supple fig. S7A and S7B). Of these, high levels of CD200 mRNA were particularly evident in dormant tumor cells from Tgfbr2^MyeKO^ mice (Figure 6A) and the elevated expression was further validated at the protein level by spectral flow studies (Figure 6B, and Supple fig. S7C). CD200 is a membrane glycoprotein structurally related to B7 family that signals through CD200R1, promoting immune tolerance and NK inactivation ^61–63^. To see if this inhibitory pathway contributed to immune evasion by the dormant tumor cells, we used shRNA to knock-down expression of CD200 in D2A1 tumor cells (Figure 6C) and observed fewer dormant lesions when these cells were injected into Tgfbr2^MyeKO^ mice (Figure 6D), consistent with a role for CD200 in dormant tumor cell resistance to immune effector function.

**Figure 6.**
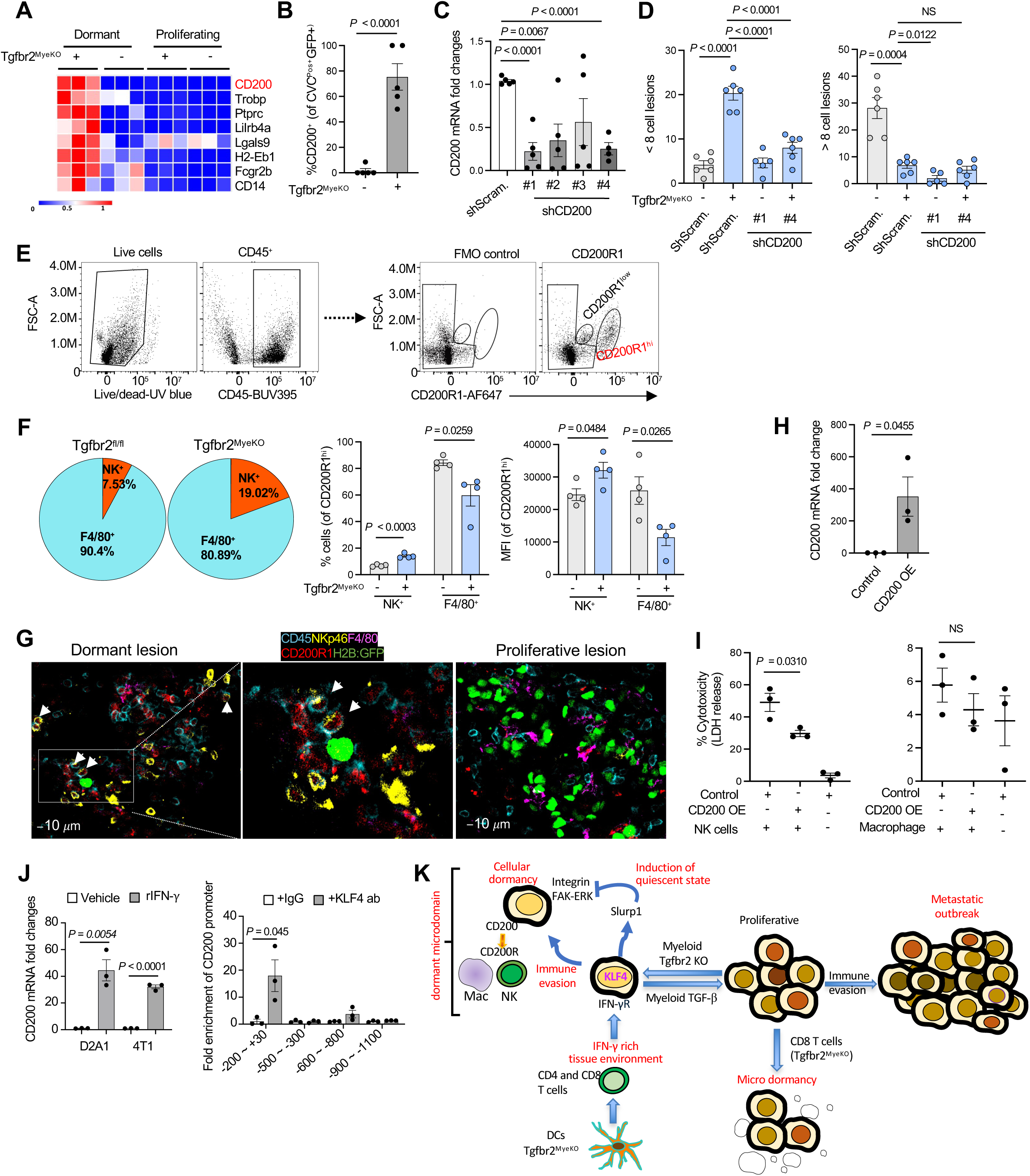
Mechanisms of immune evasion of dormant tumor cells. **A.** Heatmap of the immune evasion genes enriched from the dormant tumor cells from the Tgfbr2^MyeKO^ mice. n=3 mice per group. **B.** % CD200^+^ cells in CVC^Pos^ dormant tumor cells by Cytek analysis. n=5 mice per group. **C.** RT-qPCR for validation of CD200 knockdown in D2A1 cells. n=4 biological replicates. **D.** Number of dormant lesions (left) and total tumor lesions (right) from D2A1 cells with CD200 KD and control. n=5-6 mice per group. **E.** Flow cytometry gating of CD200R1^+^ immune cells (left), and CD200R1^Hi^ and CD200R1^Low^ subclusters (right panels). **F.** Pie chart of major immune cell subsets for CD200R1^Hi^ (left) and percentage and mean immunofluorescence intensity (right) of CD200R1^Hi^ in NK and macrophage cells comparing Tgfbr2^MyeKO^ and flox cont. mice. n=4 mice per group. **G.** Representative images of CD200R1^+^ NK and macrophages in the proximity of dormant and proliferative metastatic lesions. **H**. RT-qPCR validation of CD200 overexpression in D2A1 tumor cells. **I** % cytotoxic of CD200R1^+^ NK (left) and macrophage cells (right) co-cultured with D2A1 tumor cells with CD200 overexpression. n=3 biological replicates. **J.** left: RT**-**qPCR CD200 mRNA fold change in cultured D2A1 and 4T1 tumor cells in response to IFN-γ treatment; right: fold enrichment of CD200 expression from KLF4-ChIP qPCR of D2A1 cells with KLF4 overexpression. n = 3 biologically independent experiments. Results are shown as mean ± SD. **K.** Schematic hypothesis for immune-mediated dormancy upon abrogation of myeloid TGF-β signaling: myeloid *Tgfbr2* deletion enhances cDC function and activates CD4 and CD8 T cells leading to an IFN-γ rich tumor microenvironment. IFN-γ-induces KLF4-mediated SLURP1 expression which blocks the Integrin-FAK-ERK and induces tumor cell quiescent state. Within the dormant microdomain, the quiescent tumor cells upregulate the immune evasion program through KLF4-CD200-CD200R1 mediated inactivation of NK cells. The CD8 T cells mediated micro tumor dormancy has also been observed in Tgfbr2^MyeKO^ mice as we previously reported. All data are presented as mean ± s.e.m. *P*-values were derived from a two-tailed student t-test. Significance was determined by a *P*-value< 0.05.

To further dissect mechanism(s) of immune evasion mediated by CD200, we focused on CD200R1^Hi^ immune cells which included the NK cells and macrophages. Only NK cells but not macrophages displayed an increased cell number and CD200R1^Hi^ phenotype when comparing cells recovered from Tgfbr2^MyeKO^ with those from control mice (Figure 6E and F). A very small number of CD3 T cells were CD200R1^Hi^ but no difference was found in the percentage or MFI of these T cells between Tgfbr2^MyeKO^ and WT mice (Supple fig. S7D). To probe the potential function of CD200 in regulating NK activity, we set up an *in vitro* co-culture of CD200-overexpressing D2A1 tumor cells (Figure 6H) with CD200R1^Hi^ NK or CD200R1^Hi^ macrophages as a control. There was a clear decrease in the cytotoxicity induced by CD200R1^Hi^ NK cells (Figure 6I, left panel) but not by macrophages (Figure 6I, right panel). These data indicate one mechanism by which dormant tumor cells escape immune clearance is inhibition of NK cell-mediated cytotoxicity through CD200-CD200R1 interactions.

Connecting these data with our earlier observations on the role of IFN-γ in promoting dormancy, we found increased CD200 expression in cultured D2A1 and 4T1 tumor cells in response to IFN-γ treatment (Figure 6J, left panel). Since IFN-γ induced KLF4 expression, we looked for KLF4 binding sites in both mouse and human CD200 promoters and indeed saw the presence of the KLF4 binding motif CCCCACCC in these promoters using the Homer assay (Supple fig. S7G). Further, KLF4-ChiP assays of the CD200 promoter in D2A1 tumor cells with KLF4 overexpression revealed that KLF4 protein was enriched at -200∼+30bp region of the CD200 promoter (Figure 6J, right panel). Together, the findings suggest that myeloid *Tgfbr2* deletion enhances CD103^+^ DCs function and activates CD4^+^ and CD8^+^ T cells leading to an IFN-γ rich tumor microenvironment. IFN-γ in turn induced KLF4-mediated expression of SLURP1 that is critical in promoting tumor dormant state. These data fit with the previous conclusion that dormancy is induced by IFN-γ and these cells escape NK-mediated killing, which we now can ascribe in large measure to KLF4-mediated upregulate of an immune evasion program involving CD200-CD200R1 (Figure 6K).

### Human correlative studies of SLURP1, TβRII, and CD200-CD200R1

Are these mouse findings relevant to humans? SLURP1 expression was the highest in Basal and Her2+ forms of the four breast cancer subtypes in both METABRIC and TCGA datasets (Figure 7A). High levels of SLURP1 predicted a decreased relapse-free survival (RFS) especially for patients treated with chemotherapy (Figure 7B, left) but not those without treatment (Figure 7B, right). Additionally, among the Luminal B patients, high SLURP1 correlated with a decreased distant metastasis free survival (DMFS) for patients treated with endocrine therapy (Figure 7C, left) but not those without treatment (Figure 7C, right) (Supple fig. 8A). This inverse correlation of high SLURP1 expression with patient survival was also found in lung cancer and myeloma patients in TCGA datasets (Supple fig. 8B). Notably, SLURP1 but not GAS6, a published dormancy marker, predicted a decreased RFS in breast cancer patients treated with chemotherapy compared with those untreated (Figure 7B and C, Supple fig. 8C). These data suggest a potential role of SLURP1 in tumor dormancy and its association with treatment relapse.

**Figure 7.**
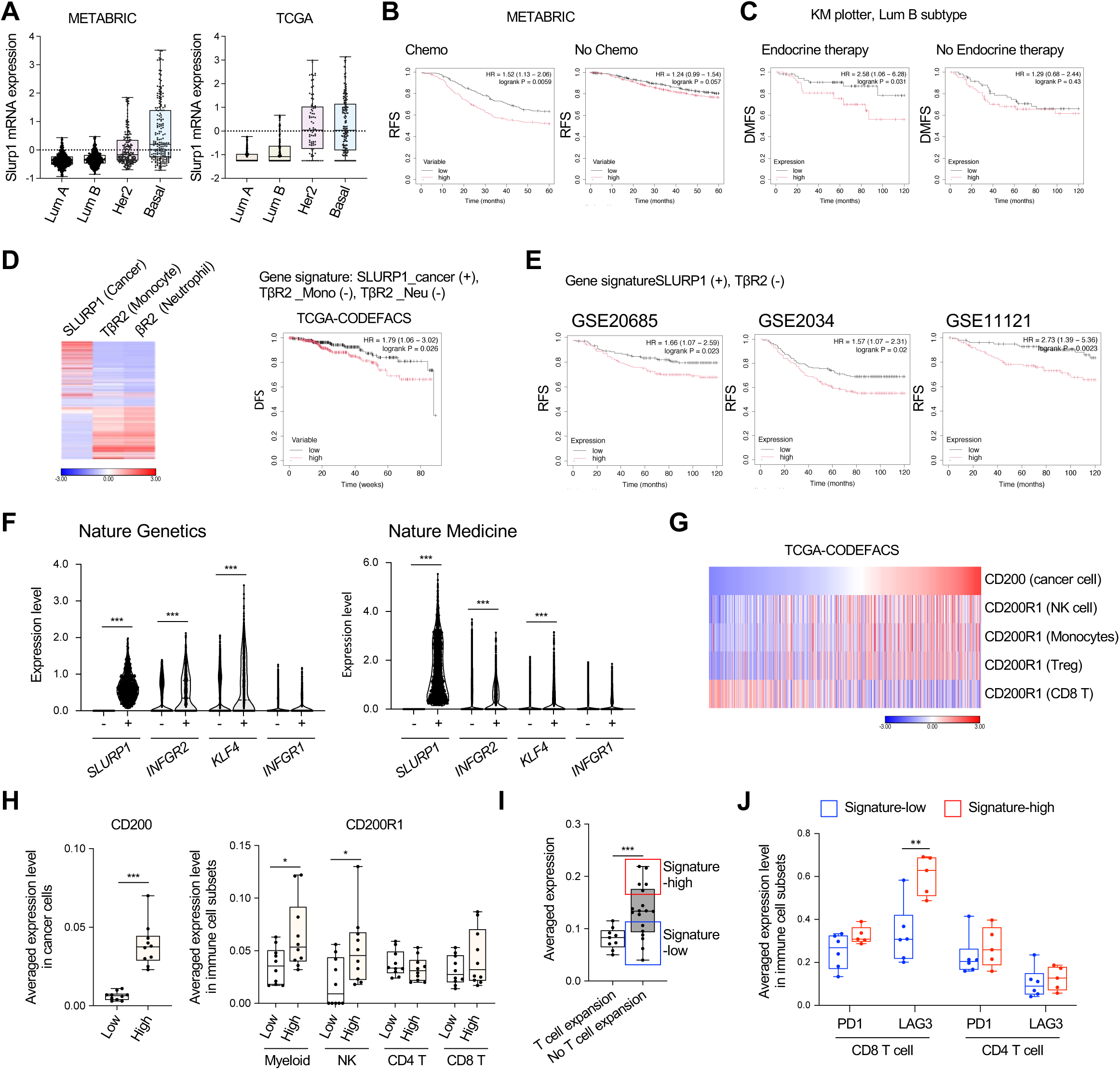
Correlative studies of SLURP1, TβRII, and CD200-CD200R1. **A.** SLURP1 expression in four subtypes of breast cancer from METABRIC and TCGA datasets. **B.** Kaplan-Meier relapse-free survival (RFS) of SLURP1 mRNA expression in breast cancer patients with (left) or without chemotherapy (right). **C.** Kaplan-Meier distant metastasis-free survival (DMFS) of SLURP1 mRNA expression in luminal B breast cancer patients with (left) or without endocrine therapy (right). **D.** Heatmap of the SLURP1expression in cancer cells and TβRII expression in monocytes and neutrophils; and their correlation with Kaplan-Meier disease-free survival (DFS), TCGA CODEFACS, deconvoluted datasets with cell type identification (https://zenodo.org/record/5790343). **E.** Correlation of SLURP1 levels in cancer cells and TβRII in monocytes and neutrophils with Kaplan-Meier RFS of patients in additional breast cancer datasets. **F.** Violin plots of SLURP1, INFGR2, KLF4, and IFNγR1 in cancer cells from single-cell RNA sequencing datasets of human breast cancer. Cancer cells were divided by SLURP1 expression (SLURP1-negative cells in blue, SLUPR1-positive cells in red). **G.** Heatmap analysis of CD200 in cancer cells, CD200R1 in CD56 NK, CD14 monocytes, Treg cells, and CD8 T cells from the TCGA-CODEFACS human breast cancer dataset. **H.** Correlation of CD200 in cancer cells with CD200R1 in myeloid, NK, CD4 and CD8 T cells. Patients were grouped by average CD200 expression in cancer cells (low in blue, high in red). **I.** Correlation of dormancy gene signature (expression levels of SLUPR1, INFGR2, KLF4, CD200 in cancer, and CD200R1 in NK cells) with status of T cell expansion in patients who received anti-PD1 treatment. **J.** The expression of PD1 and LAG3 in T cells from patients without T cell expansion, in correlation with high or low dormancy gene signature. All box plots show the 25^th^ percentile, the mean, the 75^th^ percentile and minimum/maximum whiskers. P-values were derived from a two-tailed student t-test for box and violin plots. Statistical significance was determined by log-rank test for survival analysis.

We further examined the correlation of patient survival with the dormancy regulatory mechanisms we uncovered, including increased SLURP1 in cancer cells and downregulation of TβRII in monocytes and neutrophils (Figure 7D, left). Computationally deconvolving the TCGA-BRCA dataset using CODEFACS (https://zenodo.org/record/5790343) ^64^ revealed that a signature of high Tu-SLURP1 and negatively weighted Mye-TβRII correlated with decreased survival (Figure 7D, right). There was no such correlation when survival was examined with respect to SLURP1 in cancer cells alone, or TβRII in monocytes alone, or TβRII in neutrophils alone (Supple fig. 8D). The inverse correlation of combined expression of SLURP1 and negatively weighted TGFBR2 with decreased survival was also found in other breast cancer datasets even without deconvolution or the cell identification within the TME (Figure 7E). We next looked into the correlation of SLURP1 expression with IFN-γR2 and KLF4-, the regulatory axis for SLURP1 induction, using publicly available single cell RNAseq of patient samples. The cancer cells for each patient were divided into SLURP1- or SLURP1+ subsets. We found that the expression levels of IFN-γR2 and KLF4 were significantly higher in SLURP1-positive cancer cells compared to SLURP1-negative cancer cells, whereas no significant difference was observed for IFN-γR1 in two datasets ^65, 66^ (Figure 7F).

The critical role of the CD200-CD200R1 in immune evasion of dormant tumor cells (Figure 6) led us to investigate its relevance in cancer patients. Using the deconvolved TCGA-BRCA dataset, we found that patients with high CD200 levels in cancer cells also showed high levels of CD200R1 in NK cells, CD14+ myeloid cells, and Treg (Figure 7G), consistent with our observations in mouse models. However, such high CD200R1 expression was not seen for CD8 T cells (Figure 7G), suggesting that CD200-CD200R1-mediated immune evasion is not a mechanism that affects CD8 T cell anti-tumor responses. These results were further supported by another analysis of single cell RNA-seq dataset of a human breast cancer, which revealed that patients with high CD200 expression in cancer cells showed significantly increased CD200R1 expression in NK cells and myeloid, but not in CD4 and CD8 T cells (Figure 7H).

We next generated a dormancy gene signature from our findings that included the expression levels of SLURP1, INFGR2, KLF4, and CD200 in cancer cells along with CD200R1 in NK cells and tested for links to immunotherapy responses. High expression of this dormancy signature was correlated with a lack of T cell expansion in breast cancer patients received anti-PD1 treatment (Figure 7I, Supple fig. 8E) in studies that used T cell expansion to predict immunotherapy response ^66^. Although most patients with this high dormancy signature did not show a T cell expansion response to ICB, not all patients lacking this response has a high dormancy score. We therefore further divided the patients into dormancy high- and low-signature groups (Figure 7I) to analyze immune check point genes (PD1, LAG3) or cytotoxicity genes (GZMB, GZMA, IFN-γ, TNF-α). Patients with a high dormancy signature showed significantly higher LAG3 expression in CD8 T cells, while no significant differences were observed for other genes (Figure 7J, and Supple fig. 8F). Together, these human correlative analysis from our dormancy mechanisms provide additional patient stratification and targeting strategies.

## Discussion

Our findings demonstrate that dormant tumor cells persist over long periods by co-opting conserved growth arrest and immune evasion mechanisms regulated by the innate and adaptive limbs of the immune system. Abrogation of myeloid-specific TGF-β signaling induced tumor dormancy by promoting an IFN-γ rich tissue microenvironment that activates KLF4-mediated cellular reprogramming. The IFN-γ-KLF4 axis induces increased SLURP1 expression that promotes a quiescent state through inhibition of Integrin-FAK-ERK signaling, as well as a KLF4-CD200 axis that leads to immune evasion through engagement of CD200R1^+^ on NK cells. Furthermore, our study revealed two distinct spatially localized immune niches surrounding metastatic tumor nests: Niche 1 populated with CD4^+^ and CD8^+^ T cells, alveolar macrophages, macrophages, and various DCs associated with proliferative tumor lesions and Niche 2 enriched with CD103^+^ cDCs, monocytes, neutrophils, NK cells surrounding tumor cells with the characteristics of cellular dormancy. These findings reveal critical cellular and molecular mechanisms for the induction of tumor cell quiescence and immune evasion, two of the most critical tumor dormancy hallmarks in the context of immune regulation and TME, while highlighting the complex spatiotemporal changes that lead to a mix of cellular dormancy and tumor growth control.

The tumor dormancy phenotype in our Tgfbr2^MyeKO^ mice is striking in showing that a single gene deletion in immune cells can induce tumor dormancy *in vivo*. Given the scarcity of suitable mouse models, our system advances the study of immune-regulated tumor dormancy beyond traditional paired proliferative and non-proliferative tumor cell line comparisons such as 4T1/4T07 and D2A1/D2.OR ^5, 44, 45^. It is worth noting the observed shift in dormant versus proliferative lesions in our mutant animals, despite stable total lesions counts, highlights the pivotal role of host immunity in tumor dormancy regulation (Figure 1A, 1B, 1I, 1J, Supple fig. 1F, 1G).

Many papers in the literature associate various mediators in the TGF-β pathway with regulation of tumor dormancy, including BMP, TGF-β2, and TGF-βRIII ^23, 43, 67^. Our previous studies and the present report specifically point to the critical role of myeloid cell TGF-β signaling in immune regulation of tumor dormancy. Tumor-associated myeloid cells are abundant in the tumor stroma and immune organs. These cells include neutrophils, tumor-associated macrophages (TAMs), dendritic cells (DCs), and monocytic and granulocytic myeloid-derived suppressor cells (MDSCs). The latter in particular contribute to compromised host immune surveillance and cancer recurrence ^68, 69, 8, 70–77^. Our previous studies demonstrate that myeloid TGF-β signaling is critical in immune/inflammatory homeostasis as its abrogation inhibited tumor metastatic progression and its prolonged absence also induced spontaneous stroke in aged mice ^47–49, 59^. Together with the results presented here, we propose that myeloid TGF-β signaling acts as a switch: the “ON” mode promotes immune suppression and metastatic outgrowth, while the “OFF” mode fosters anti-tumor immunity and tumor dormancy.

Multiple immune cells play roles in dormant niche formation and tumor cell emergence from dormancy, including neutrophils ^21, 78, 79^, macrophages ^76, 77^, DCs ^12^, CD4 T ^80^ and CD8 T cells, and NK cells ^12, 29, 36, 38, 39, 81, 82^. However, the mechanism underlying cellular dormancy regulated by host immunity *in vivo* is particularly under-studied. Our work revealed that cellular dormancy is associated with localized regions enriched in NK cells, CD103^+^ cDCs, monocytes, and neutrophils. These regions are distinct from those surrounding proliferative tumor lesions that have abundant CD4^+^ and CD8^+^ T cells, alveolar macrophages, and DCs. *In vivo* depletion of cDC and CD8 T cells established causal roles of these cells in the observed cellular tumor dormancy. The immune niche for dormant lesions is associated with IFN-γ production upon abrogation of myeloid TGF-β signaling ^47–49^, a conclusion supported by a prior report ^83^ and a study using single RNA-seq ^82^.

It was surprising given these findings that our imaging studies showed that the CD103 cDC and the CD8 cells involved in IFN-γ production and induction of cellular dormancy were not immediately adjacent to the quiescent tumor cells, which resided in Niche 2 with colocalized NK cells. One possible explanation for these findings is that the adaptive immune response takes time to develop after tumor introduction. The early arriving tumor cells would enter the proliferative phase and their antigens would promote the adaptive response, but by the time the activated CD4 and CD8 T cells arrive at the metastatic locale, the growing tumor cells are not readily induced into the dormant state and are controlled by active anti-tumor effector function in Niche 1. Other tumor cells that are delayed in proliferation and that likely express high levels of the IFN-γR can sense the IFN-γ produced by these effector lymphocytes. Several studies measuring the dimensionality of IFN-γ action in tissues show that there are biologically meaningful effects 100-300 um from the producer cells ^84–86^. This is more than sufficient to act outside of the definition we have for Niche 1 radius, maintaining the quiescent state of tumor cells capable of SLURP1-mediated dormancy induction. CD200 expression that is induced at the same time would then protect these cells from killing by the NK cells in the Niche 2 environment. Detailed studies that combine spatial transcriptomics and highly multiplex protein imaging will be needed across time series to test this hypothesis.

The immune evasion mechanism involving CD200-CD200R1 mediated inactivation of NK cells in the dormant niche adds to the existing body of work showing cellular dormancy is related to evasion of CD8^+^ T cell– and NK cell-mediated detection and clearance of the tumor cells ^21, 36, 37, 39^. These insights may provide an explanation for post-ICB relapse in which the effective adaptive anti-tumor responses are dependent on cDC1 and CD8 T cells, with this same response leading to the development of immune-resistant dormant tumor cells by IFN-γ. The latter are presumably responsible for late arising metastatic lesions in patients with apparent remissions.

Our study identified SLURP induction by IFN-γ induced signaling as a key step in induction of cellular dormancy. Functional and biochemical experiments suggest that SLURP1 functions through a cell-matrix interface by interacting with fibronectin-integrin pathways to block FAK-mediated cell proliferation. This is supported by previous reports of downregulation of uPAR under these conditions that lead to reduced integrin and MAPK signaling ^51^. Extracellular matrix has been reported to play an important role in both sustaining dormancy and in tumor cell emergence from this state. For example, dormant tumor cells upregulate COL17A1 or type III collagen, which to contribute to maintenance of dormancy ^31, 32^. Chemotherapy disrupts COL17A1 and inhibits the dormant state through FAK-YAP activation ^31^. On the other hand, degrading the fibronectin matrix by matrix metalloproteinase 2 (MMP-2) promotes cancer cell outgrowth ^30^. Additionally, the interaction of L1CAM with laminin in perivascular basement membranes also promotes exit from dormancy ^87^. In part this is mediated through the activation of YAP and MRTF ^81^. Targeting integrin αvβ6-TGFβ-SOX4 pathway by an integrin αvβ6/8-blocking monoclonal antibody sensitizes TNBC cells to cytotoxic T cells in highly metastatic murine TNBC models that were poorly responsive to PD-1 blockade ^88^. A key challenge moving forward is identifying the SLURP1 interacting extracellular proteins and elucidating their functional roles in dormancy regulation.

Our studies highlight the KLF4-SLURP1-Integrin-FAK-ERK and CD200-CD200R1 axes as vulnerabilities of dormant cell states and the molecular determinants of communication with their residing niches. The strong correlation between these pathways and breast cancer patient survival, treatment relapse, and immune inhibitory mediators such as PD-1 and LAG3 highlights their potential as therapeutic targets for preventing and treating metastatic diseases. Future direction can be focused on exploring the therapeutic potential of targeting CD200 and LAG3 in preclinical mouse models with an inducible myeloid-Tgfbr2 knockdown system, facilitating the translation of dormancy-related insights into therapeutic applications. This is particularly important in addressing challenges associated with ICB and dormancy-associated relapse. Notably, in early-stage breast cancer patients treated with endocrine therapy, metastases can emerge gradually between 5 to 20 years post-diagnosis and treatment ^3, 11, 89, 90^. Drugs that directly kill the dormant cells or maintain them in a dormant state will likely provide long-term benefits for breast cancer patients ^25, 39, 91–93^, as supported by an NR2F1 agonist that suppresses metastasis by inducing cancer cell dormancy ^26, 27^. However, the subtle interplay of immune factors in mediating both tumor cell control versus induction of dormancy that preserves viable cells able to emerge as late clinically relevant metastases suggests that harnessing this information effectively will require exceedingly careful attention to effective balancing the desired vs. the undesired effects of immune manipulation.

## Supporting information

Yang Supple fig and table

## Acknowledgments

We thank Dr. William Telford for his support in characterizing the immune cells. We acknowledge CCR Sequencing Facility and bioinformatics team at the Frederick National Laboratory for Cancer Research (FNLCR) for library preparation, sequencing, and primary QC analysis. We thank Drs. Xuan Qi and Maxwell Lee for their effort in analyzing the immunofluorescence images. We are grateful for technical assistance from the CCR animal facility. This work was supported by NCI intramural funding to Dr. Li Yang, and was supported in part by the DIR, NIAID, NIH [Lymphocyte Biology Section] and by a cooperative agreement between CCR, NCI and DIR, NIAID [Center for Advanced Tissue Imaging].

## Author contributions

A. Ahad designed, planned, and performed most of the experiments, analyzed data, and contributed to the intellectual progression of the project and manuscript draft. F. Leng contributed to the initiation and intellectual progression of the project, mouse model characterization, and dormant tumor cell validation. H. Ichise, E. Schrom, and R. Germain lead the immune niche characterization, including experimental planning, data analysis, results interpretation, and manuscript preparation. J.Y. So performed human correlative analysis. C. Sellner generated IFN-γR KD, CD200 OE, and CD200 KD cell lines, performed Western, and contributed to manuscript preparation. Y. Gu and W. Wang performed ChIP and PCR assays. C. Lieu and F. Livak supported the flow imaging analysis. K. Wolcott supported the dormant cell sorting. W.Y. Park and R. Yang were involved in cell line generation for *in vivo* mouse model experiments. M. Kruhlak supported the CD200R1 imaging experiment. O. Aprelikova, J. Gray, and V. Kopardé established the dormant cell sorting, subsequent RNA-seq, and its analysis. Y. Moriwaki provided the SLURP1 antibody and contributed to critical reading of the manuscript. L.Yang initiated, organized, and designed the study, and supervised the overall project and the manuscript preparation.

## Material and Methods

### Mouse strains and cell lines

*Tgfbr2* floxed (Flox cont.) mice were bred with *Lysozyme 2* promoter-driven *Cre* recombinase (LysM-Cre) mice to generate mice with myeloid-specific deletion of *Tgfbr2* (Tgfbr2^MyeKO^) as previously described ^1^. Transgenic mice expressing *Tgfbr2*-shRNA were crossed with rtTA floxed mice (Strain #005670, The Jackson Laboratory) and then bred with LysM-Cre mice to generate Doxycycline (dox)-inducible myeloid-specific Tgfbr2 knock-down (Tgfbr2^MyeKD^) mice. The Tgfbr2^MyeKO^ and Tgfbr2^MyeKD^ mice genotypes were confirmed by PCR analysis of tail biopsies, using primer sequences listed in Supplementary Table 1. Female mice aged 6–8 weeks were used for animal studies. All animal procedures reported in this study were performed by NCI-CCR affiliated staff following the animal protocol LCBG007-4 and were approved by the NCI Animal Care and Use Committee (ACUC), with federal regulatory requirements and standards. AAALAC International accredits all components of the intramural NIH ACU program. All mice were maintained at a 12 h light-dark cycle, 64 – 72 °F (18 – 22 °C) temperature, and 30 – 70 % humidity condition and euthanized by following the NCI CO2 euthanasia protocol.

4T1 (#CRL-2539) cells were purchased from the American Type Culture Collection, D2.OR and D2A1 cell lines were kindly provided by Dr. Ann Chamber’s lab. The TSAE1 cell line was kindly provided by Dr. Lalage M. Wakefield of the Laboratory of Cancer Biology and Genetics, National Cancer Institute. Lenti-X 293T cells (#632180), a subclone of HEK 293 cells, were purchased from Takara Bio. All cell lines were cultured in DMEM medium (Gibco) supplemented with 10% FBS (Gibco), 100 U/ml penicillin, and 50 μg/ml streptomycin (Sigma, St Louis). All cell lines were confirmed to be mycoplasma negative by the MycoAlertTM Mycoplasma Detection Kit (Lonza) and kept in liquid nitrogen when not used.

### Mouse models of metastasis and tumor dormancy

For orthotopic metastasis studies, 4T1 cells (2×10^5^) were injected into mice’s mammary fat pad (MFP) #2. For experimental metastasis models, 4T1 cells (1×10^5^), D2A1 cells (2×10^5^ - 1×10^6^), or TSAE1+mHer2 cells (5×10^5^) were injected through the tail vein (TVI). The number of lung metastatic nodules was evaluated by Indian ink staining by direct counting via a dissection microscope or by lung surface imaging.

For dormancy-associated studies, when indicated, 4T1 or D2A1 cells were stained with CellVue Claret (CVC) dye (Millipore Sigma, MINICLARET) according to the manufacturer’s instructions followed by TVI. For lung surface imaging, 1mg/ml of doxycycline (Millipore Sigma, #D9891) was added to the mice water, or Cumate (Millipore Sigma, # 268402) dissolved in 70% ethanol (Sigma, # 268402) was subcutaneously injected 3 days before endpoint to induce the expression of H2B-GFP. After the animals were sacrificed, lungs were inflated with PBS, imaged using the EVOS Cell Imagining System (Thermo Fisher, M7000), and H2B-GFP+ tumor cells were individually counted. 1-8 celled H2B-GFP+ lung lesions were considered dormant lesions and lesions with greater than 8 cells were considered proliferating lesions.

### In vivo immune cell depletions

All immune cell depletion antibodies were purchased from BioXcell. For in vivo depletion of CD103^+^ DCs, mice were intraperitoneally injected with InVivoMAb anti-mouse CD103 (M290; 150ug/mouse) or IgG2a isotype control (2A3; 150ug/mouse) antibodies every 3 days starting from day -3 of D2A1 cell injection until mice were sacrificed on day 12. For pDCs depletion, mice were intraperitoneally injected with InVivoMAb anti-mouse CD317 (927; 150ug/mouse) or IgG2b isotype control (LTF-2; 150ug/mouse) antibodies every 3 days starting from day -3 of D2A1 cell injection until mice were sacrificed on day 12. For CD8^+^ T cell depletion, mice were intraperitoneally injected with *InVivo*MAb anti-mouse CD8α (YTS 169.4, 200ug/mouse) or IgG2b isotype control (LTF-2; 200ug/mouse) antibodies every 3 days starting from day -1 of D2A1 cell injection until mice were sacrificed on day 12.

### Single-cell suspension

Lungs were collected from tumor-bearing mice, minced on ice, and transferred to gentleMACS C Tubes (Miltenyi, 130-093-237) containing RPMI (Gibco) and tumor digestion enzymes (Miltenyi, #130-093-237). Lung tissue was digested using the gentleMACS™ Dissociator (Miltenyi,130-096-730) according to the manufacturer’s instructions. Following tissue digestion, red blood cells were lysed with ACK lysis buffer (Quality Biological, # 118-156-101) and cells were resuspended with RPMI supplemented with 10% FBS. The cells were washed with PBS and filtered through a 70uM cell strainer to generate a single-cell suspension for downstream analysis.

### Imaging flow cytometry

CVC labeled H2B-GFP D2A1 and 4T1 cells were injected through the tail vein, and the lungs were collected 12-14 days after injection. 3-5 million single suspended cells isolated from lung tissue were stained with 1:1000 LIVE/DEAD™ Fixable Near IR Dead Cell Stain Kit (Thermo Fisher, L34975) for 30 min at room temperature. Cellular non-specific binding sites were blocked with Fc blocking antibody (CD16/32 antibody, BioLegend, #101320) for 15 min. The cellular samples were then fixed, permeabilized, and stained with conjugated with P-p38-PE (Invitrogen, #12-9078-42), GAS6 (R&D systems, #AF986), and SLURP1(abbexa, #abx129448) antibodies for 40 minutes. After washing, the samples were stained with secondary antibody BV421 (dilution ratio) for Slurp1 and AF568 (dilution ratio) for GAS6 for 30 min. All the staining incubations were performed at 4 ℃ in dark conditions. The fluorescence intensities were measured with a 2-camera, 4-laser, 12-channel Amnis ImageStream MkII (Cytek Bioscience) imaging flow cytometer. The images were generated with INSPIRE software and further analyzed for fluorescence intensities on IDEAS 6.02 (Cytek Bioscience).

### Cell sorting

CVC labeled H2B-GFP D2A1 and 4T1 cells were injected through the tail vein and the lungs were collected 12-14 days after injection. Dissociated cells were sorted on a FACSAria IIu (BD Biosciences) cell sorter equipped with 405nm violet, 488nm blue, and 642nm red lasers, using 100um nozzle low-pressure conditions. Single color controls were used for spectral compensation and to establish sorting gates to collect GFP+ CVC^Pos^ and GFP+ CVC^Neg^ cells.

### Immune cell profiling with spectral flow cytometry

A multicolor antibody panel covering myeloid to lymphoid cells was used to analyze cells on the Aurora spectral flow cytometer (Cytek Bioscience). The list of antibodies is provided in Supplementary Table 2. 3-5 million single suspended cells from lung tissue were blocked with Fc blocking antibody (anti-CD16/32 antibody, BioLegend, #101320) for 30 min at 4 ℃ and stained with 1:1000 LIVE/DEAD™ Fixable Blue Dead Cell Stain Kit (Thermo Fisher, L34962). Next, cells were stained with the corresponding antibody cocktail at 1:100 dilution prepared in FACS buffer (PBS with 2% FBS and 1mM EDTA) for 30 min at 4 ℃. Following 30 min cell fixation (Thermo Fisher, # 00-5523-00), cells were permeabilized and intracellularly stained with intracellular antibodies at 1:60 dilution for 40 min at room temperature. Samples were acquired on a Cytek Aurora spectral flow cytometer (Cytek Biosciences), and data were analyzed in FlowJo software v10.6.0 (BD Bioscience).

To examine effect of SLURP1 on downstream integrin signaling, D2A1 cells were cultured in Matrigel and treated with 50nM rSLURP1 (Abbexa, abx166123) or rGST (Abbexa, #abx655551) as a control for 24 hour. Cells were collected from Matrigel culture as previously described with a few modifications (PMC8202162). In brief, cells were washed with warm PBS and incubated for 30 min in chilled cell recovery solution (Corning, # 354253). Cells were then treated with TrypLE Express Enzyme (Gibco, # 12604-021) for 10 min at 37°C. Cells were fixed with fixation buffer (Invitrogen, # 00-5523-00) and washed with FACs buffer to make single cell suspension for staining.

### *Ex-vivo* co-culture of tumor cells and immune cells

Flox cont. and Tgfbr2^MyeKO^ mice were tail vein injected with D2A1 cells. 12 days later, CD8^+^ T cells and CD103^+^ DCs were sorted from the lungs. Immune cells were cultured with CVC-labeled H2B-GFP-D2A1 cells in GlutaMAX RPMI 1640 medium (Gilbco, #61870036), supplemented with 10% FBS, 1000 IU/ml murine recombinant IL-2 (Peprotech, #212-12), 1ug/mL anti-CD3 (eBioscience, # 16-0031-82), and 1ug/mL anti-CD28 (eBioscience, #14-0281-82) in the presence or absence of IFN-γ (BioXcell, BE00550) or TNF-α (BioXcell, BE0058) neutralization antibodies for 3-5 days. 24 hours before Ki-67 and Cleaved caspase3 staining, cells were treated with 2μg/ml dox to induce H2B-GFP expression in D2A1 cells. The ratio of D2A1 cells to CD103^+^ cDCs (1:1) and CD8+ T cells (1:2) was used.

To examine the effect of CD200 cancer cell expression on NK cell and macrophage cytotoxicity, flox cont. and Tgfbr2^MyeKO^ mice were tail vein injected with D2A1 cells. NK cells and macrophages were sorted from CD200R1^+^ splenocytes 12 days later and cultured overnight in RPMI 1640 (Gibco) with 10% FBS (Gibco), 0.375% sodium bicarbonate, 1mM sodium pyruvate, 0.0001% 2-mercaptoethanol and 1000 IU/ml murine recombinant IL-2. Control or CD200 overexpression D2A1 cells were then added to immune cell cultures for 24 hours before measuring cytotoxicity using the CytoTox96 Cytotoxicity Assay (Promega, # G1780). The ratio of D2A1 cells to NK cells and macrophages was 1:1. Percent cytotoxicity was calculated using a maximum LDH release control and the formula: *Precent cytotoxicity* = 100 × 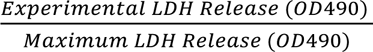 as described via manufacturer instructions.

### Mouse tissue immunofluorescence staining using iterative bleaching and extended multiplexity (IBEX) and analysis

Lung tissues were prepared, imaged, and aligned as previously described with minor modifications (https://www.biorxiv.org/content/10.1101/2024.07.04.601620v1). In brief, lungs were inflated with Cytofix (BD) and kept in BD Cytofix/Cytoperm (BD BioScience) at 4 °C for 14-16h. The next day, tissues were washed with PBS once, incubated in 30% sucrose for 24h at 4 °C, embedded in OCT compound (Tissue-Tek), and kept at -80℃ until use. Frozen lung tissues were cut at 20-30 μm on a CM3050S cryostat (Leica) and adhered to Superfrost Plus slides (VWR, Cat. # 48311-703) coated with 15 μL of chrome alum-gelatin adhesive (Newcomer Supply, Cat. #1033A). 20-30 μm lung tissue serial sections were adhered to chrome alum gelatin (Newcomer Supply) coated slides and kept overnight to dry. Tissue sections were permeabilized in BD perm buffer supplemented with mouse Fc block (BD) and Hoechst 33342 (Biotinum). Serial sections were screened with a Leica THUNDER imaging system equipped with a 20⨉ objective (NA 0.75) (Leica, Cat #11506343), and dormant tumor cells/colonies (less than 8 cells) were identified based on GFP signals and used for subsequent IBEX procedures. Tissues were stained with PELCO BioWave Pro-36500-230 microwave (Ted Pella) for staining. Two cycles of 2–1–2–1–2–1–2–1–2 program were used for immunolabeling with the BioWave, where “2” denotes 2 min at 100 W and “1” denotes 1 min at 0 W. After staining, slides were mounted with Fluormount G (Southern Biotech, Cat. #0100-01) and examined on a Leica THUNDER Imaging system or STELLARIS confocal microscope. Fluorochrome inactivation was performed with 1 mg/mL of LiBH_4_ (STREM Chemicals) for 15 minutes. Samples were further stained with the antibodies for the subsequent cycle of imaging. SimpleITK program was used for image alignment, followed by downstream analysis. Confocal images were collected using a Leica Stellaris 8 laser scanning microscope equipped with an HC plan-apochromat 20× (N.A. 0.75) multi-immersion objective lens with 0.284 μm x-y pixel size and 4.0 um optical section thickness. The antibodies were provided in Supplementary Table 3.

### IBEX analysis

#### Preparing Multiplex Mouse Lung Images for Analysis

Each channel of the mouse lung multiplex images was restricted to its dynamic range of expression by saturating the brightest 0.01% of pixels. Images were then converted to 8-bit format. Noise due to autofluorescence and channel spillover was evident by its similar appearance across multiple channels, and it was reduced via pairwise channel arithmetic. The Hoechst nuclear stain was binarized to segment individual cells, followed by a watershed step to separate touching nuclei. Raw signal for every membrane marker (all markers except Ki67, GFP, and αGFP) was passed through a Gaussian filter with a four-pixel radius to increase the overlap of membrane stain with the nucleus of the expressing cell. The size, centroid location, and mean fluorescence intensity (MFI) of each marker was then measured for each cell in each image. All steps were performed in ImageJ.

#### Classifying Cells into Phenotypes

Using the single-cell MFIs for each marker, a threshold was drawn at the “elbow” of the expression distribution for each marker to separate positive from negative expression. Each cell’s profile of +/- expression for each marker was compared to expected expression profiles for known cell types, and cells that matched an expected type were categorized as such. The MFIs for these cells were then used to define representative quantitative expression profiles for each known cell type.

The quantitative expression profile for each remaining uncategorized cell was then used as the output variable in multivariate linear regression, with the representative quantitative expression profiles for each known cell type as multivariate predictors. Essentially, this approach models each uncategorized cell as a mixture of the representative expression profiles for known types. Among known types that contribute significantly to an uncategorized cell’s expression profile, that with the largest contribution was selected as the uncategorized cell’s type.

This left 7.7% of cells that were not significantly associated with a known type. These were subject to unsupervised k-means clustering to search for unexpected cell types. However, the clusters that emerged were typically characterized by the expression of just a single marker, which was visually confirmed to correspond to leftover noise in the images. Thus, uncategorized cells were excluded from the analysis. Furthermore, the 0.4% of cells categorized as γδ T cells were also excluded from analysis due to the remaining spillover of CD11b into the γδTCR channel, which made the distinction of true γδ T cells challenging. All steps were performed in R.

### Definition of Tumor Lesions

The table of XY centroid coordinates for each cell was used to measure each tumor cell’s distance from every other tumor cell, within each image. These distances defined a graph representation of the tumor cells, in which tumor cells were considered neighbors if their distance was ≤ 200 µm. Connected components on this graph then defined individual lesions.

After the tumor cells were partitioned into separate lesions, the table of XY centroids was again used to measure the distance of each non-tumor cell to each tumor cell. All non-tumor cells within 200 µm of a tumor cell were assigned to that tumor cell’s lesion. Thus, while each tumor cell was assigned to exactly one lesion, non-tumor cells could be assigned to zero, one, or more than one lesion. All steps were performed in R.

### Analyses of Whole Lungs and Tumor Lesions

The total fraction of tumor cells in Tgfrb2^MyeKo^ vs. flox cont. lungs was compared using a binomial linear regression model. The output variable was each cell’s binary status as a tumor cell or not, and the predictors were the fixed effect of genotype and the random effect of sample ID. The size of lesions in Tgfrb2^MyeKo^ vs. flox cont. lungs were compared using a zero-truncated negative binomial model. The output variable was each lesion’s size (number of tumor cells), and the predictors were the fixed effect of genotype and the random effect of sample ID. Both models were built using the glmmTMB R package.

To explore the immune cells in each lesion, the counts of each immune cell type in each lesion were re-expressed as center-log ratios (CLRs) to account for the compositional nature of this data. The CLRs for each cell type were then standardized separately to account for the differences in total abundance across cell types. These standardized CLRs were used in place of raw cell counts for all ensuing analyses (except direct ratios of CD8 and CD4 T cells).

The pairwise correlations across cell types were measured, and the modules of co-occurring cell types were defined using the gplots R package. Principal component analysis (PCA) was also performed on the standardized CLRs in R. The significance of associations between PC1 and the size and genotype of each lesion was investigated using a Gaussian linear regression model. The output variable was each lesion’s PC1 score, and the predictors were the fixed effects of lesion size, genotype, and their interaction, and the random effect of sample ID.

### Clearing enhanced 3D imaging (Ce3D)

Volumetric imaging with optically cleared samples with Ce3D was performed as described previously with slight modification (https://www.pnas.org/doi/10.1073/pnas.1708981114) (https://www.biorxiv.org/content/10.1101/2024.07.04.601620v1). Briefly, frozen lung samples were sectioned at 300 μm on a CM3050S cryostat (Leica Biosystems). The samples were hydrated and washed with PBS to remove OCT in a 24-well plate. Samples were incubated for at least 12 hours in BD Perm/Wash Buffer (BD Bioscience) containing 1% mouse Fc block (BD Bioscience, Cat. #553142) and stained with titrated antibodies in BD Perm/Wash Buffer (BD Bioscience, Cat. #554723) for 24 hours at room temperature on a shaker. Stained samples were washed with BD Perm/Wash Buffer three times for at least 20 minutes at room temperature on a shaker and transferred on a slide with two silicon isolators (Grace BioLabs, Cat. #664407) and treated with 200 μL of Ce3D medium [1.82 g Histodenz (Millipore Sigma, Cat. #D2158-100G, 0.1% triton, and 0.5% thioglycerol (Millipore Sigma, Cat. #M1753) per 1 mL 40% N-methylacetamide (Millipore Sigma, Cat. #M26305–500G) in PBS] inside a chemical fume hood and sealed with a cover slip (Electron Microscopy Sciences, Cat. #63766-01). Samples were protected from light and incubated at room temperature on a shaker overnight. Cleared samples were mounted with 40 μL of new Ce3D and sealed with a coverslip with two SecureSeal Imaging Spacers (Grace Bio-Labs, Cat. #654002) and examined on a Leica STELLARIS confocal microscope equipped with a 20 ⨉ objective (NA 0.75).

### Sphere formation assay

Matrigel (Corning, #356231) was added to 96-well plates or 15 μm-slide (ibidi, #81506) on ice and solidified by incubating at 37°C incubator for 40 min. Tumor cells were suspended in culture media with 2% Matrigel. 500 cells/ 96 wells were added with varying concentrations of rSLURP1 (Abbexa, abx166123) or rGST (Abbexa, #abx655551) as a control. Cells were cultured for 3-5 days and supplemented with fresh recombinant protein every 24 h. For Integrin activation, 1000 D2A1 cells were seeded in Matrigel-coated 96-wells. Cells were then pretreated for 1 h with 500 μg/mL integrin activation antibodies or IgG (BioXcell, BE0232, or BE0089) before adding 50nM of rSLURP1. Cells were supplemented with fresh recombinant protein every 24 h and cultured for 5 d before imaging. For rSLURP1 withdrawal, D2A1 cells in Matrigel-coated 96-wells were treated with 50nM rSLURP1 for 5 days. On day 5, cells were washed with 0% FBS media to remove recombinant protein, supplemented with fresh 10% FBS media, and allowed to grow for 3 more days before imaging. Relative proliferation and spheroid area were measured using ImageJ software (https://imagej.net/ij/).

### RNA-seq of sorted dormant, proliferative tumor cells, tumor cells with rSLURP1 treatment

GFP+ CVC^Pos^ (dormant) or GFP+ CVC^Neg^ (proliferating) D2A1 cells were sorted from the lungs of WT or Tgfbr2^MyeKO^ mice 12 days after TVI. Cells were sorted into lysis buffer and an aliquot of 50 cells (determined by Flow Sorter). For 3D spheroid cultured cells. rGST or rSLURP1 treated cells were harvested as described (PMC8202162) on day 5. Samples were subjected to Reverse Transcription using QIAGEN RNeasy Kit (#74104) followed by library preparation using Smart seq2 protocol (Nature Protocols, 9:171-181, (2014).

The sequencing reads were processed for adapter trimming using Cutadapt (version 1.14) to remove unwanted adapter sequences and low-quality bases at the read termini. The sequencing quality of the trimmed reads was assessed per sample using FastQC (version 0.11.5) to evaluate base quality scores, adapter contamination, and sequence duplication levels. Besides FastQC, Preseq (version 2.0.3) was used to estimate library complexity and predict the yield of unique reads for deeper sequencing. Picard tools (version 1.119) were employed to assess metrics, such as insert size distribution, duplication rates, and sequence alignment summaries. To further ensure data quality, RSeQC (version 2.6.4) provided insights into RNA integrity, GC content, and potential biases in sequencing coverage.

The trimmed reads were then aligned to the mm10 mouse genome using STAR (version 2.5.2b) in two-pass mode to enhance mapping sensitivity and accuracy. After successful alignment, gene expression levels were quantified using RSEM (version 1.3.0), which employs an expectation-maximization algorithm to accurately estimate gene and transcript abundance. The reference genome annotation used for expression quantification was GENCODE annotation M12.

Gene-level expression estimates from RSEM were used in downstream DESeq2 analysis to identify differentially expressed genes between the conditions shown in Fig 2B. The statistical significance of differentially expressed genes was determined using a q-value threshold of ≤0.05 and an absolute fold change cutoff of ≥ 2. Each of these comparisons aimed to elucidate the transcriptional differences between dormant and proliferative states, as well as the impact of Tgfbr2 deletion. Further downstream analyses, including hierarchical clustering and principal component analysis (PCA), were performed to visualize sample relationships and validate the robustness of expression changes. To gain insights into the functional relevance of differentially expressed genes, pre-ranked Gene Set Enrichment Analysis (GSEA) was performed to assess pathway-level enrichment.

### Lentivirus production and transduction

The predesigned shRNA plasmids were purchased from Millipore Sigma. The catalog numbers and sequences of shRNA used are provided in Supplementary Table 4. The mRuby-p27K^-^ quiescence reporter was kindly provided by Dr. Steven Cappell from the Laboratory of Cancer Biology and Genetics, National Cancer Institute. The packaging plasmid psPAX2 (#12260) and the PMG2.G (#12259) envelope plasmid were obtained from Addgene. The mouse coding sequences for Slurp1, Klf4, and CD200 were cloned into the FerH-NeoR-Lenti overexpression vector and sourced from GenScript Biotech. For lentivirus production, 80% confluent LentiX cells were transfected with lentiviral vector, psPAX2, and PMD2.G by Lipofectamine 3000 reagent (Invitrogen, # L3000001) in OptiMem medium. The medium was replaced with fresh media 4-6 h after transfection. After 48 h, viral supernatant was collected, centrifuged, and filtered through 0.45 μm filters and stored at -80°C. For lentivirus transduction, indicated cells were incubated with viral supernatants and 8 ug/mL Polybrene for 24 h. Successfully transduced cells were selected by adding the appropriate antibiotic into the culture medium 48 -72 h after transduction.

### Slurp1 sgRNA-induced gene knockout

D2A1 cells were transduced with Pspcas9-2A-puro-px459 (Addgene 48139) constructs containing Slurp1 single-guide RNAs designed by F. Zhang’s laboratory, which was followed by 2 μg/ml puromycin selection for 2 d. Single clones were collected and knockout was confirmed by genotyping PCR fragments consisting of the target sequences. Slurp1 sgRNAs and PCR validation primer sequences are listed in Supplementary Table 5.

### RT-qPCR

RNA was isolated using the Quick-DNA/RNA Miniprep Kit (Zymo Research, # D7001), and 1μg of cDNA was synthesized using the ReverTra Ace qPCR RT Kit & Master Mix (Toyobo, # FSQ-101). RT-qPCR reactions were carried out using the PowerSYBR Green PCR Master Mix (Thermo Fisher, # 436765) on a QuantStudio 6 Flex Real-Time PCR System (Thermo Fisher, #4485691). For SLURP1 induction, D2A1 and 4T1 cells were serum starved overnight before treatment with recombinant murine IFN-γ (10ng/mL) (Cell Signaling, #39127S) for 24 hours. mRNA levels were normalized to the housekeeping gene Gapdh, and relative fold change was determined using the 2−ΔΔCt method. All primer sequences are provided in Supplementary Table 6.

### Western blotting

Cells were lysed with RIPA Lysis Buffer (Sigma, #R0278-50ML) supplemented with cOmplete protease inhibitor cocktail (Roche, #04693132001) and phosphatase inhibitor cocktail (Roche, #4906845001). Protein concentration was measured by Pierce’s BCA Protein Assay (Thermo Fisher, #23225). An equal amount of total protein was denatured, separated on NuPAGE 4-12% Bis-Tris protein gels (Thermo Fisher, #NP0321), and transferred to a nitrocellulose membrane (Bio-Rad, #1704270). Membranes were blocked in 5% non-fat milk in TBS and .1% Tween 20 (TBS-T) for 1 hour before overnight incubation in primary antibodies against SLURP1 (Provided by Yasuhiro Moriwaki, Keio University), IFNGR1 (Cell Signaling, #84318S), pSTAT1 (Cell Signaling, #9167S), STAT1 (Cell Signaling, #9172S), KLF4 (abcam, #ab129473), or β-actin (Santa Cruz, sc-69879, AC-15). Primary antibodies were diluted 1:1000. Secondary HRP-conjugated anti-mouse (Cell Signaling, #7076S) and anti-rabbit (Cell Signaling, #7074S) were used at a 1:2000 dilution. Protein detection was done using the ECL reagent (Bio-Rad, #1705061) and visualized using the ChemiDoc imaging system (BioRad).

### BioPlex Immunoassay

Flox cont. and Tgfbr2^MyeKO^ mice were tail vein injected with 2×10^5^ D2A1 cells, and lungs were collected after 12 days. Lungs were flash-frozen and stored in -80 until use. Lung tissue was homogenized with the Precellys Lysing Kit (KT03961-1-002.2, Precellys) using the Precellys 24 tissue homogenizer (P002391-P24T0-A.0, Bertin Instrument). Protein concentration of cytokines was measured and quantified using the Bio-Plex Pro Mouse Cytokine 23-plex Assay according to the manufacturer’s protocol (Biorad, M60009RDPD).

### KLF4 ChIP-Slurp1 RT-qPCR

The ChIP assay was performed following the protocol provided by Active Motif (catalog #53008). Briefly, 1 × 10⁷ KLF4 OE and control cells were treated with 1% formaldehyde for 10 min at 37°C to cross-link proteins with DNA. Cells were then lysed in SDS buffer for 10 min on ice, followed by sonication (30% of maximum power, 15 s on, 1 min rest, for a total of 25 min) to generate DNA fragments ranging from 200 to 1000 bp. The samples were precleared with protein A beads for 30 min on a rotor at 4°C and then incubated overnight at 4°C with 2 µg anti-KLF4 or control antibodies. After a series of washes, the eluted fractions were subjected to DNA isolation and analyzed by qPCR. Primer sequences used to map Slurp1 promoter regions are listed in Supplementary Table 7.

### DNase I sensitivity assay

For the DNase I sensitivity assay, KLF4 OE and control cells were collected, and nuclei were isolated following the protocol of the nuclei isolation kit (catalog #78833). The nuclei pellets were resuspended in DNase digestion buffer, and 0 or 2 U of DNase I (EN0521) was added, followed by a 5 min incubation at 37°C. After terminating the digestion, genomic DNA was extracted, and the indicated chromatin regions were analyzed by qPCR.

### Human correlative studies

Publicly available datasets from breast cancer patients (METABRIC and TCGA) were analyzed to determine *SLUPR1* expression in different subtypes. For survival analysis, patient survivals were correlated with *SLUPR1* expression or a two-gene signature combining *SLURP1* expression with negatively weighed *TGFBR2* expression in the METABRIC, TCGA and Kaplan-Meier Plotter datasets. For cell type-specific gene expression analysis, the computationally deconvolved TCGA-BRCA dataset by CODEFACS (https://zenodo.org/record/5790343) was utilized (ref). To define a cell type-specific gene signature, *SLURP1* expression from cancer cells was integrated with negatively weighed *TGFBR2* expression from monocytes and neutrophils. To assess the correlation between SLUPR1 and dormancy-related genes, cancer cell-specific expression data of *SLURP1*, *INFGR2*, *KLF4*, and *INFGR2* were extracted from two scRNA-seq datasets: Nature Genetics (2021) (https://singlecell.broadinstitute.org/single_cell/study/SCP1039/a-single-cell-and-spatially-resolved-atlas-of-human-breast-cancers) and Nature Medicine (2021) (https://lambrechtslab.sites.vib.be/en/single-cell). To investigate CD200-CD200R1 correlation, as well as the link between dormancy-related genes and immunotherapy response, cell annotation, patient metadata, and cell type-specific gene expression were extracted from the Nature Medicine (2021) sgRNA-seq data.

### Statistics & Reproducibility

All data are presented as mean ± Standard Error of the Mean (s.e.m.). Unless otherwise indicated, comparisons between two groups were performed using a two-tailed Student’s t-test. Statistical analysis of survival data used the Wilcoxon test. Statistical analyses were performed using GraphPad Prism 7.0 software, and significance was defined as *P* <0.05. Significance is noted in figure or figure legends; *P*-value as ^∗^*P* < 0.05, ^∗∗^*P* < 0.01, ^∗∗∗^*P* < 0.001, ^∗∗∗∗^*P* < 0.0001, NS > 0.05. No data were excluded from the analyses, except for Supplementary Figure 1A, in which one replicate was excluded due to an outlier. All mice used in *in vivo* studies were randomized before cancer cell injection. The phenotype of animal experiments was not evaluated blindly, and multiple researchers verified major results.

## Data availability

The data supporting the findings of this study are available within the article, and the supplementary information files are available from the corresponding author upon request. The raw and processed RNA-sequencing data have been deposited in the GEO database and are publicly available under accession numbers (GSE292496) and (GSE292514). All relevant source data for each figure are provided.

