## Supplementary material for "Immune Niche Formation in Engineered Mouse Models Reveals Mechanisms of Tumor Dormancy": Yang Supple fig and table

Supplemental Fig. 1

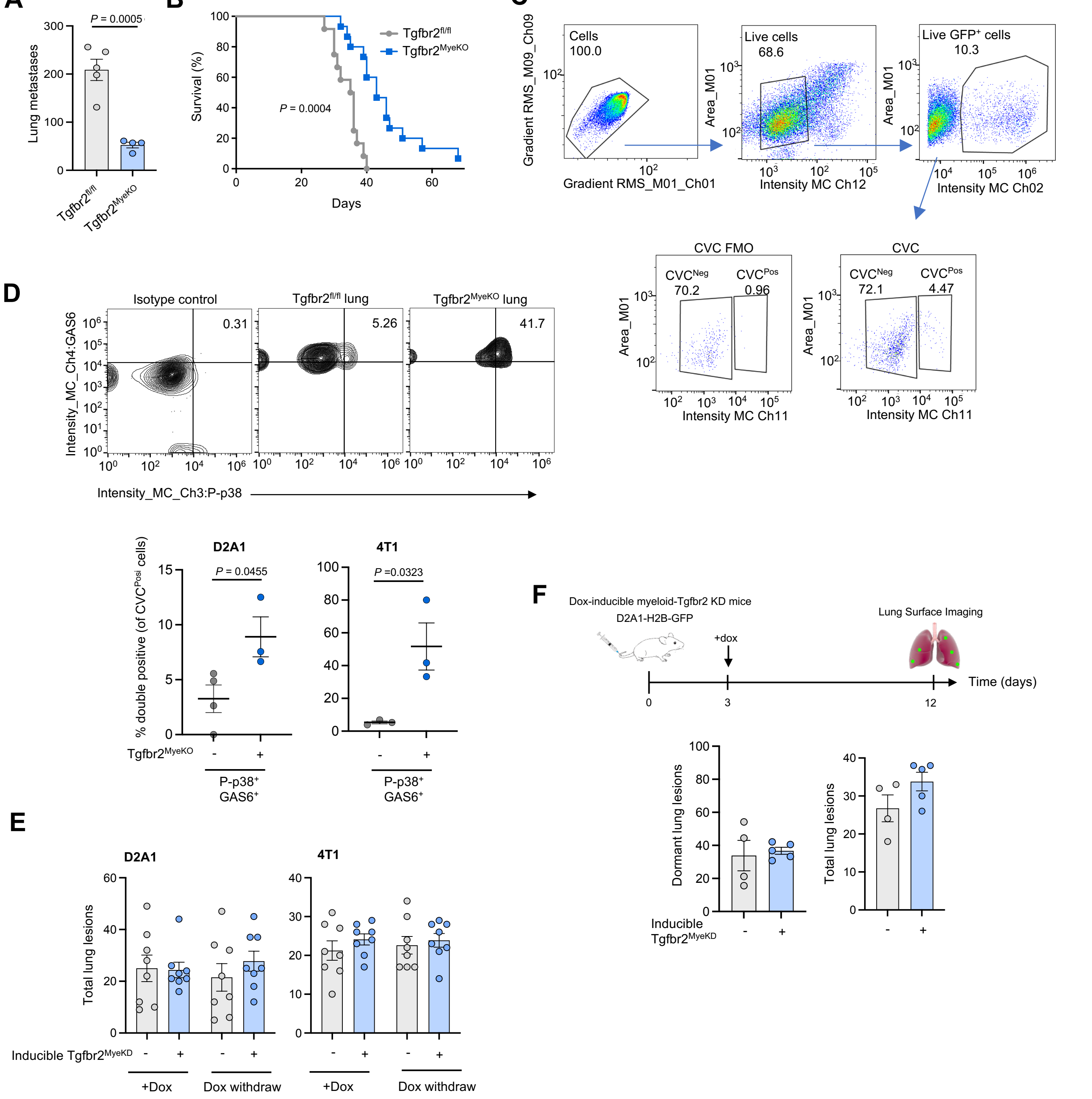


**Figure S1. Tumor dormancy induction mediated by abrogation of myeloid-specific TGF-β signaling. A.** Decreased lung metastasis in Tgfbr2$^{MyeKO}$ mice compared to flox cont. mice that received tail vein injection (TVI) of D2A1 cells (n=4-5 mice per group). **B.** Tgfbr2$^{MyeKO}$ mice survived longer than flox cont. after receiving TVI of D2A1 cells. Statistical analysis by Wilcoxon test. **C.** Imaging flow cytometry gating strategy for GFP$^{+}$/CVC$^{Pos}$ D2A1 cells. **D.** Imaging flow gating strategy for P-p38 and GAS6 (left) and graphs showing more CVC$^{Pos}$ D2A1 and 4T1 cells were P-p38 and GAS6 double positive from Tgfbr2$^{MyeKO}$ mice than flox cont (n=3-4 mice per group). **E.** Total lung lesions did not change with myeloid Tgfbr2 KD or re-expression in the D2A1 TVI model (left) at and 4T1 orthotopic model (right) (n=8 mice per group). **F.** Dox-induced myeloid-Tgfbr2 KD after D2A1 TVI did not display a tumor dormancy phenotype suggesting the importance of a preestablished lung microenvironment (n=4-5 mice per group). Statistical analysis by unpaired two-tailed t-test. All error bars represent mean $\pm$ s.e.m.

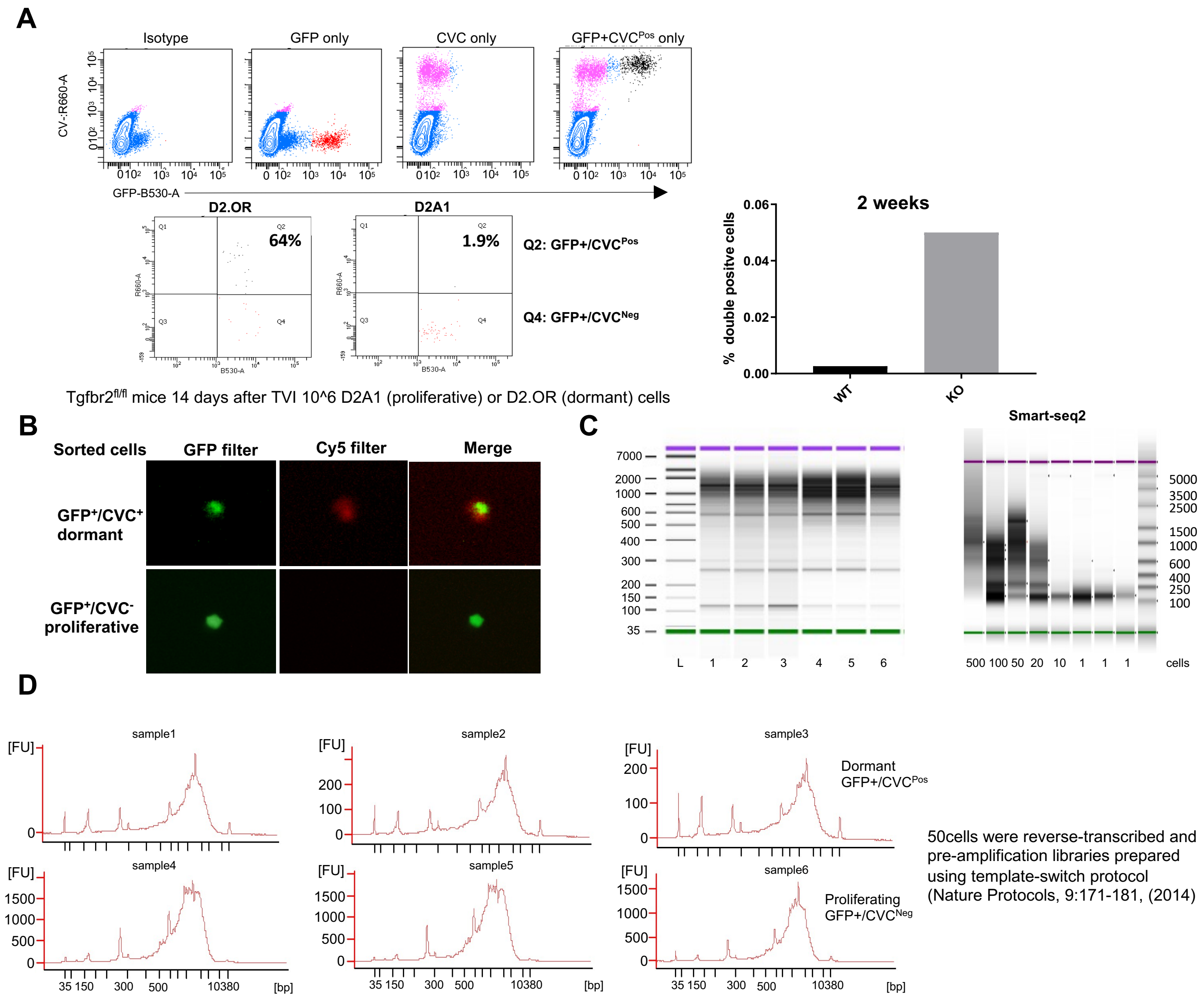


**Figure S2. Identification of dormant tumor cells and RNA sequencing validation A.** Flow cytometry plots for sorting dormant (GFP+CVC$^{Pos}$) vs proliferative (GFP+CVC$^{Neg}$) cancer cells before TVI (top panel) and flow cytometry gating of GFP+CVC$^{Pos}$ D2.OR and D2A1 lung metastasis (bottom left panel) and quantification showing more GFP+CVC$^{Pos}$ double positive D2A1 lung metastasis from Tgfbr2$^{MyeKO}$ mice compared to flox cont. **B.** Sorting validation of GFP+CVC$^{Pos}$ and GFP+CVC$^{Neg}$ D2A1 cells **C.** RNA quality for sequencing. **D.** Pre-amplification libraries from dormant cells (n=50) after reverse transcription.

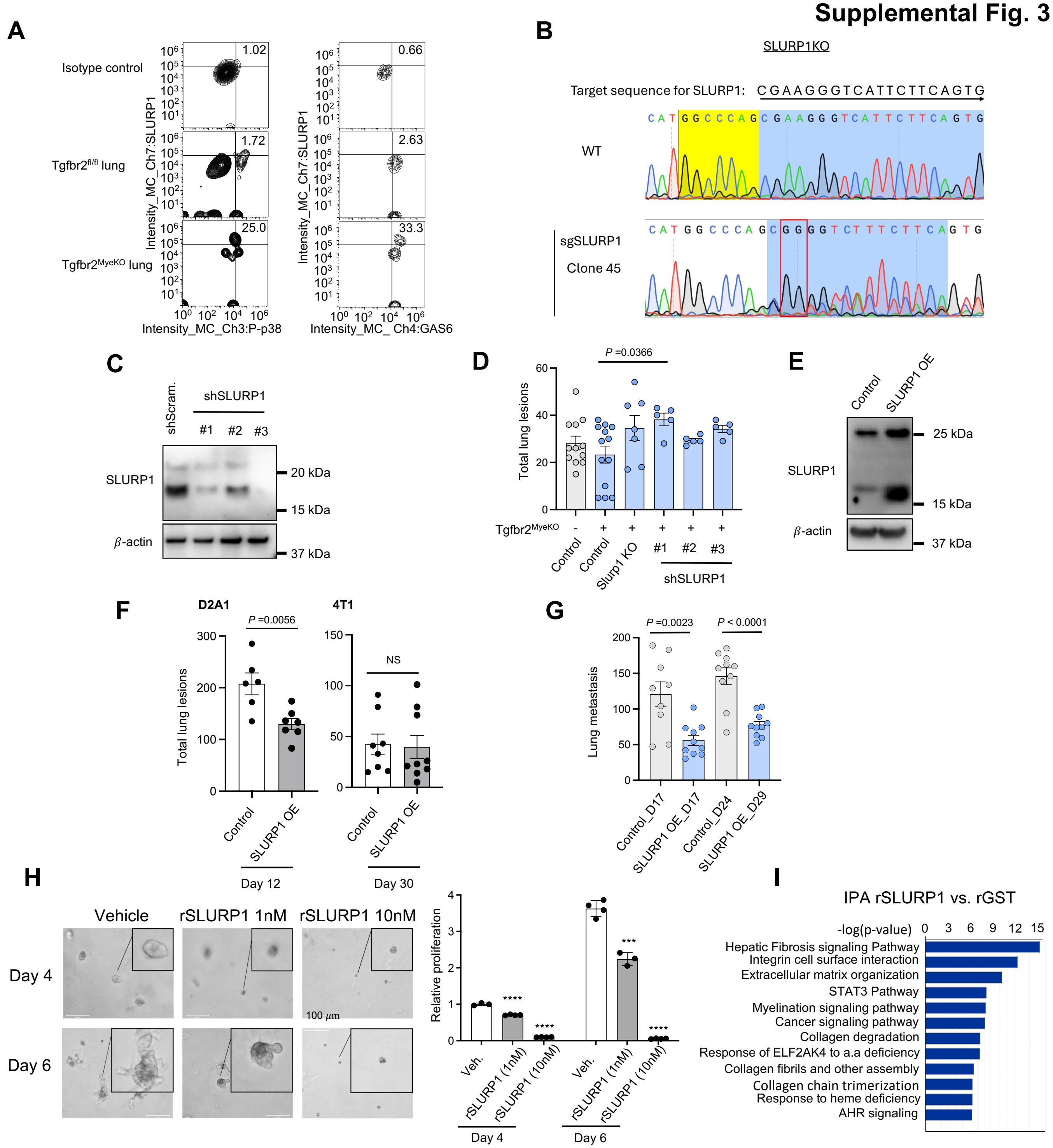


**Figure S3. SLURP1 regulation of tumor cell proliferation and dormancy. A.** Imaging flow cytometry gating for the expression of SLURP1, P-p38, and GAS6 in dormant (GFP+CVC$^{Pos}$) and proliferative (GFP+ CVC$^{Neg}$ ) D2A1 cells. **B.** Sanger sequencing chromatograms for Slurp1 KO clone. **C.** Western blot confirming KD of SLURP1 in D2A1 cells. **D.** Total tumor lesions mostly did not differ between Slurp1 KO and KD and control D2A1 cells in Tgfbr2$^{MyeKO}$ mice (n=5-13 mice per group). **E.** Western blot of SLURP1 overexpression in 4T1 cells. **F.** Total lung lesions from D2A1 (left) and 4T1 cells (right) with SLURP1 overexpression **G.** Metastasis nodule count by Indian Ink staining from mice that received TVI of SLURP1 overexpressing D2A1 cells (n=9-10 mice per group). **H.** Recombinant SLURP1 inhibited D2A1 spheroid growth in a dose-dependent manner (left) and quantitative data (right) (n=3-4 biological replicates).**p<0.01, ***p<0.001, ****p<0.0001 **I.** IPA of rGST vs. rSLURP1 treated D2A1 cells in 3D culture. Statistical analysis by unpaired two-tailed t-test. All error bars represent mean ± s.e.m.

### Supplemental Fig. 4

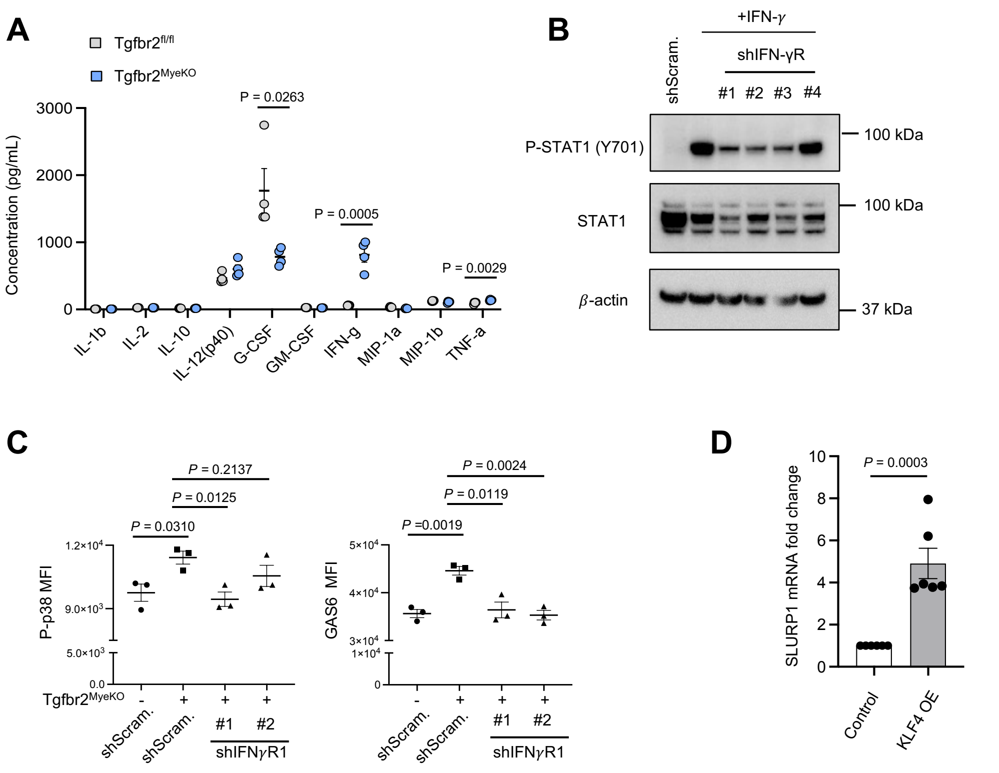


**Figure S4. Mechanisms of SLURP1 regulation A.** Bio-Plex Immunoassay showing increased IFN-$\gamma$ and TNF-$\alpha$ in the lung lysate from tumor-bearing Tgfbr2$^{MyeKO}$ mice (n=4 mice per group). **B.** Western blot confirming reduced STAT1 phosphorylation in shRNA IFN-γR KD cells following treatment with recombinant IFN-$\gamma$ (100ng/mL) for 30 minutes. **C.** P-p38 and GAS6 expression from Imaging flow cytometry of dormant cells with or without IFN-γR KD, single cell suspension from the lungs of the tumor-bearing mice. n=3 biologically independent experiments. **D.** RT-qPCR showing over-expression of KLF4 increases SLURP1 mRNA in D2A1 cells (n=6 biological replicates). Statistical analysis by unpaired two-tailed t-test. All error bars represent mean ± s.e.m.

### Supplemental Fig. 5

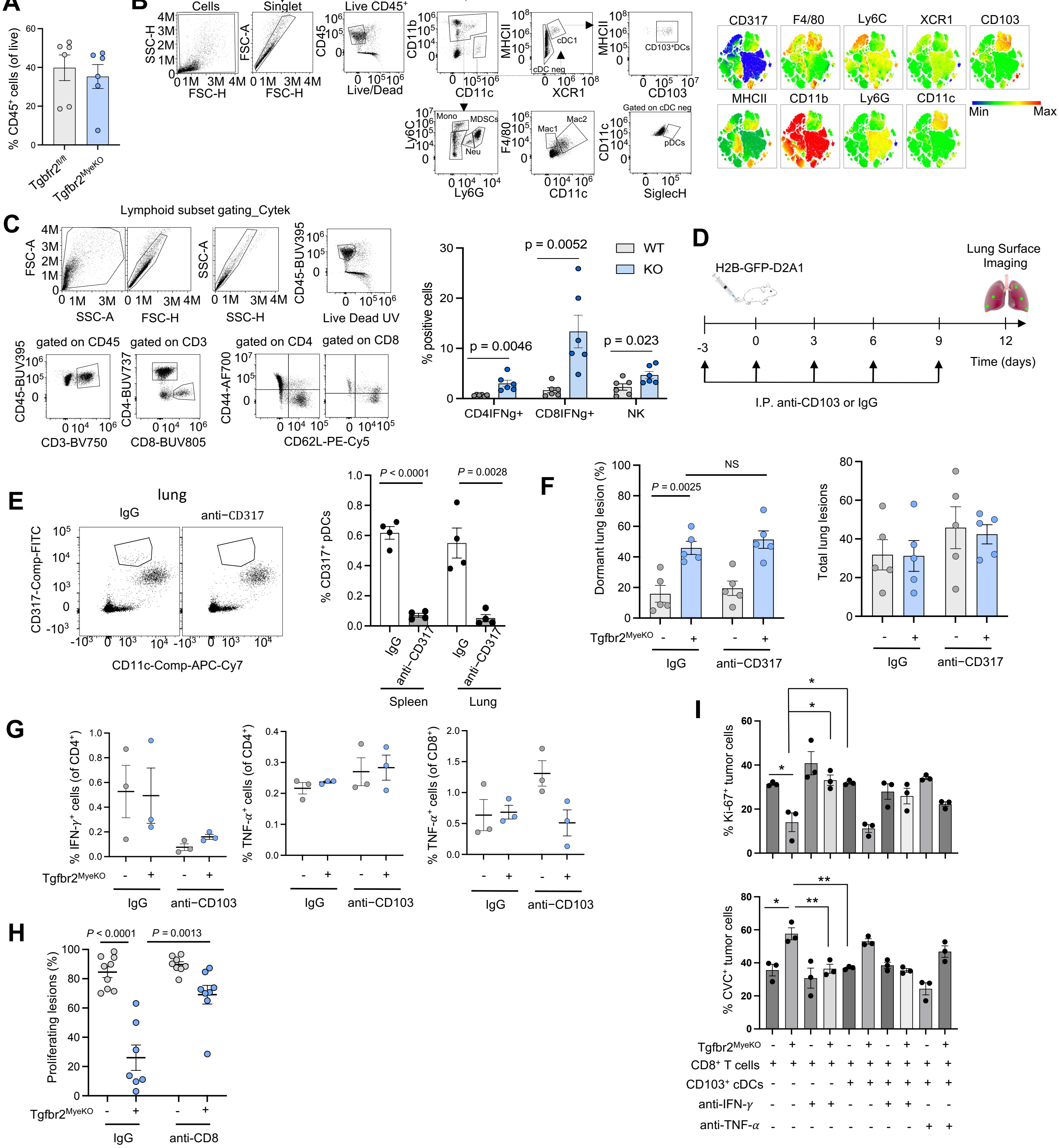


**Figure S5. IFN-γ mediated immune surveillance in tumor dormancy. A.** Cytek of the CD45+ cells from lungs of Tgfbr2MyeKO and flox cont. mice. **B**. Cytek gating strategy and tSNE plots for myeloid cell subclusters; the expression intensity for each marker is indicated by the color scale bar. **C.** Cytek gating strategy for lymphoid cell subsets (left). And % IFN-$\gamma$+ T cells and NK cells (right). **D.** Experimental design for the depletion of CD103+ cDCs. **E.** Gating strategy (left) and validation of CD317+ pDCs depletion in spleens and lungs of mice (right) (n=4 mice per group). **F.** No difference in dormant lung lesions upon CD317+ pDC depletion (n=5 mice per group). **G.** Flow cytometry analysis of IFN-$\gamma$ and TNF-$\alpha$ expression in CD4+ T cells and TNF-$\alpha$ expression in CD8+ T cells upon CD103+ cDC depletion (n=3 mice per group). **H.** CD8+ T cell depletion increased proliferative lesions. **I.** Bar plot showing the percent of Ki-67+ (top) and CVCPos (bottom) D2A1 cells in the presence of CD8+ T cells or CD103+ cDCs or IFN-$\gamma$ or TNF-$\alpha$ neutralizing antibodies (n=3 biological replicates). *p<0.5, **p<0.01. Statistical analysis by unpaired two-tailed t-test. All error bars represent mean ± s.e.m.

Supplemental Fig. 6

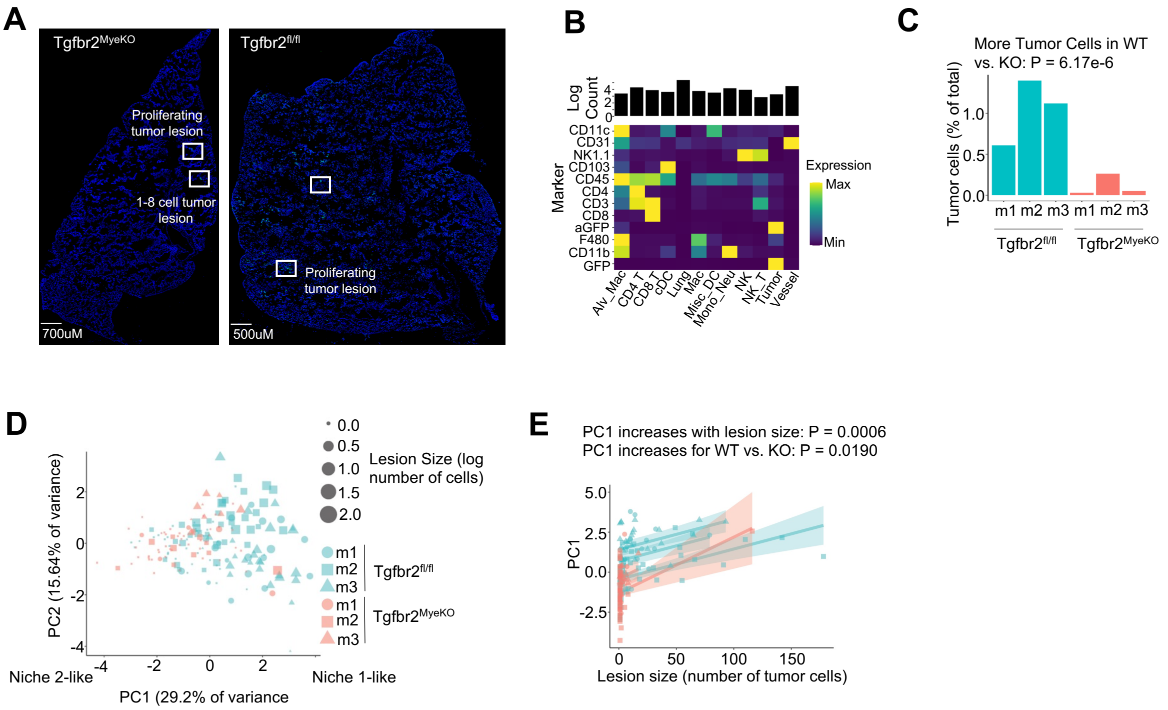


**Figure S6. Characterization of immune niches for dormant and proliferative tumor lesions A.** Whole lung tissue with representative imaging of tumor lesions. **B**. Clustering showing average cell markers profiles for the 12 known cell types identified from mouse lung IBEX images. **C**. Percentage of tumor cells in each mouse lung sample. Flox cont. mice contain more tumor cells (binomial regression model, P = 6.17 × 10-6). **D.** PCA of tumor lesions based on standardized CLRs for each immune cell type**. E.** PC1 scores for each tumor lesion as a function of lesion size and genotype. Smaller lesions and lesions from Tgfbr2$^{MyeKO}$ mice have lower PC1 scores on average (Gaussian regression model, P = 0.0006 and P = 0.0190, respectively).

**Supplemental Fig. 7**

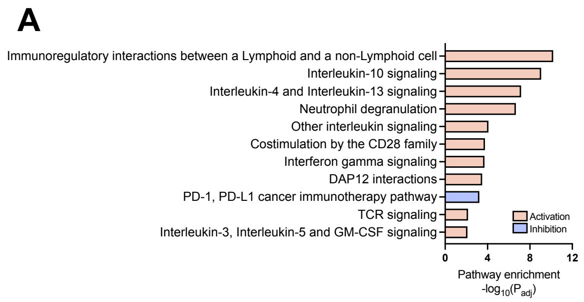


**B**

| Pathway | Genes |
|---|---|
| Immunoregulatory interactions between a Lymphoid and a non-Lymphoid cell | c3,cd200,cd200r1,cd300c,cd300ld,cd300lf,cd3e,cd3g,cd40,cd8a, colec12,crtam,fcgr2b,fcgr3a/fcgr3b,hcst,itgb2,klrg1,lilra5,lilrb3, lilrb4, ncr1,pilrb,sftpd,sh2d1b,slamf6,trem1,trem2,tyrobp |
| Interleukin-10 signaling | ccl2,ccl4,ccl5,ccr1,ccr2,ccr5,cd86,fcer2,il10ra,il1rn,ptafr,ptgs2,tnf |
| Interleukin-4 and Interleukin-13 signaling | batf, ccl2, f13a1, fcer2, hgf, ighg, il13ra2, itgam, itgax, itgb2, ndn, osm, pik3r1, ptgs2, rora, s1pr1, tnf |

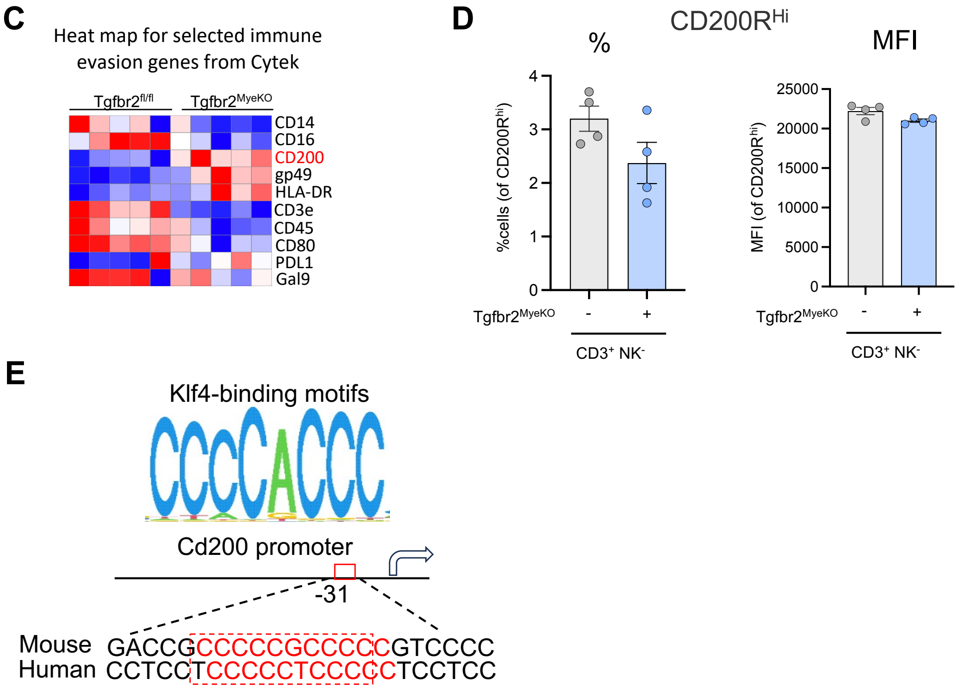


**Figure S7. Immune evasion of dormant tumor cells. A.** Pathway analysis from RNA-seq data comparing dormant vs. proliferative tumor cells indicating potential immune evasion mechanisms in tumor dormancy. **B.** Top gene candidates from the top three differential pathways and identification of CD200-CD200R1 regulatory axis. **C.** Heat map from Cytek analysis for CD200 validation among other immune modulatory genes. **D.** no difference in % $CD3^{+}CD200R^{Hi}$ T cells nor in $CD200R^{Hi}$ MFI comparing $Tgfbr2^{MyeKO}$ and flox cont. mice (n=4 mice per group). **E.** KLF4 binding site mapping in the *Cd200* promoter by Homer assay. KLF4 binding motif CCCCACCC was shown in mouse and human *CD200* promoters. Statistical analysis by unpaired two-tailed t-test. All error bars represent mean ± s.e.m.

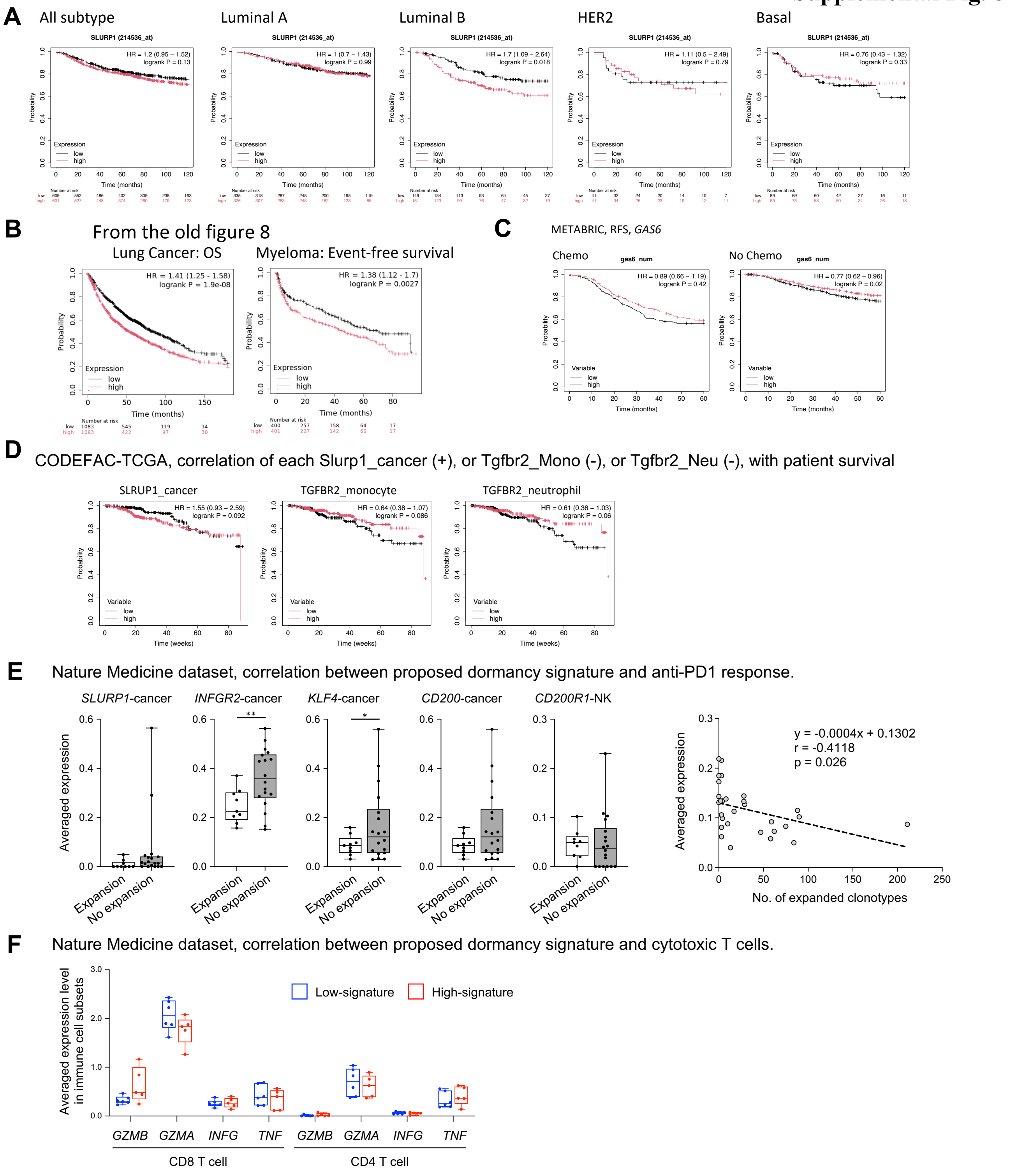


**Figure S8. Human correlative studies of SLURP1, TβRII, and CD200-CD200R. A.** High SLURP1 predicted a decreased DMFS for Luminal B patients (Earlier 2017, Exclude biased dataset) Q1 vs Q3, by KM-plotter. **B.** High SLURP1 predicted a decreased OS or Event free survival for lung cancer and myeloma patients, TCGA dataset. **C.** SLURP1 but not GAS6 predicted an increased RFS in breast cancer patients treated with chemotherapy compared with those untreated. **D.** The correlation of SLURP1 in cancer cells alone, or TβRII in monocytes alone, or TβRII in neutrophiles alone with patient survival in breast cancer patient dataset from TCGA CODEFACS, **E.** Average expression levels of *SLURP1*, *INGFR2*, *KLF4*, *CD200* from cancer cells and *CD200R1* from NK cells comparing patients with and without T cell clonotype expansion (left 5 panels). Correlation between dormancy gene signature and number of expanded clonotypes for each patient (right panel). **F.** Average expression levels of *GZMB*, *GZMA*, *INFγ*, *TNF* in CD8 and CD4 T cells comparing patients with low and high dormancy signatures.

Supplementary Table 1. Primers for mouse genotyping.

| Mouse Strain | Primer | Sequence |
|---|---|---|
| LysM-Cre | LysM-Cre-R | GTTGCATCGACCGGTAATGCA |
| | LysM-common-F | CTTGGGCTGCCAGAATTTCTC |
| | LysM-wt-R | TTACAGTCGGCCAGGCTGAC |
| Tgfbr2-floxed | Tgfbr2-flox-F | TAAACAAGGTCCGGAGCCCA |
| | Tgfbr2-flox-R | ACTTCTGCAAGAGGTCCCCT |
| rtTA-floxed | rtTA-F | TGCCGCCATTATTACGACAAGC |
| | rtTA-R | ACCGTACTCGTCAATTCCAAGGG |
| Tgfbr2-shRNA | Tgfbr2-shR-F | CCATGAAGATCAAGGTGGTCGA |
| | Tgfbr2-shR-R | CCGTCTTCGTATGTGGTGATTCT |

### Supplementary Table 2. The list of anti-mouse antibodies used in CyTEK12

| Antigen | Cat # | Clones | Company |
|---|---|---|---|
| CD45 | 564279 | 30-F11 | BD biosciences |
| CD11b | 612977 | M1/70 | BD biosciences |
| CD11c | 117310 | N418 | Biolegend |
| CD206 | 141714 | C068C2 | Biolegend |
| F4/80 | 749283 | T45-2342 | BD biosciences |
| Ly-6G | 127629 | 1A8 | Biolegend |
| Ly6C | 128036 | HK1.4 | Biolegend |
| PDL1 | 563369 | MIH5 | BD biosciences |
| iNOS | 12-5920-82 | CXNFT | eBioscience |
| Arg1 | 17-3697-82 | A1exF5 | eBioscience |
| CD103 | 121408 | 2E7 | Biolegend |
| CD200 | 565547 | OX-90 | BD biosciences |
| CD200R | 566345 | OX-110 | BD biosciences |
| B220 | 751580 | RA3-6B2 | BD biosciences |
| CD40 | 124618 | 3/23 | Biolegend |
| CD44 | 103056 | IM7 | Biolegend |
| CD62L | 104410 | MEL-14 | Biolegend |
| CD3 | 100249 | 17A2 | Biolegend |
| CD4 | 553043 | RM4-5 | BD biosciences |
| CD25 | 102004 | PC61 | Biolegend |
| CD8a | 612898 | 53-6.7 | BD biosciences |
| FoXP3 | 126406 | MF-14 | Biolegend |
| IFNg | 505830 | XMG1.2 | Biolegend |
| IL-6 | 561376 | MP5-20F3 | BD biosciences |
| TNFa | 506338 | MP6-XT22 | Biolegend |
| Tbet | 561263 | O4-46 | BD biosciences |
| TIM3 | 134012 | B8.2C12 | Biolegend |
| PD1 | 109112 | RMP1-30 | Biolegend |
| NK1.1 | 560618 | PK136 | BD biosciences |
| Perforin | 154306 | S16009A | Biolegend |
| TCR γ/δ | 118124 | GL3 | Biolegend |
| IL-2 | 503824 | JES6-5H4 | Biolegend |
| IL-17A | 506927 | TC11-18H10.1 | Biolegend |
| IL-2 | 503824 | JES6-5H4 | Biolegend |
| IL-13 | 159403 | W17010B | Biolegend |
| IL-10 | 563277 | JES5-16E3 | BD biosciences |
| Granzyme B | 515406 | GB11 | Biolegend |
| CD80 | 46-0801-82 | 16-10A1 | Bioscience |
| CD86 | 105027 | GL-1 | Biolegend |
| Ki67 | 556027 | B56 | BD biosciences |
| CD317 | 127105 | 129C1 | Biolegend |
| pFAK (Y397) | ab81298 | EP2160Y | abcam |
| pERK (T202, Y204) | 560115 | 20A | BD biosciences |
| p-p38 (T180, Y182) | 12-9078-42 | 4NIT4KK | Invitrogen |

### Supplementary Table 3. The list of anti-mouse Antibodies using in IBEX

| Antigen | Cat # | Clones | Company |
|---|---|---|---|
| CD45 | 58-0451-82 | 30-F11 | Invitrogen |
| CD11b | 101217 | M1/70 | Biolegend |
| F4/80 | 41-4801-82 | | Invitrogen |
| Ly6G | 46-9668-82 | 1A8 | Invitrogen |
| CD11c | 117346 | | biolegend |
| CD3 | 100282 | 17A2 | Biolegend |
| CD8 | 100708 | 53-6.7 | Biolegend |
| CD103 | MAB1990-SP | 262523 | R&D |
| CD200R | 566345 | OX-110 | BD biosciences |
| Cl-Caspase3 | D3E9 | | CST |
| Ki67 | 48-5698-82 | | Invitrogen |
| FoXP3 | 50-5773-82 | FJK-16s | Invitrogen |
| NFAT | (D43B1) XP ® | | CST |
| CD4 | 41-0042-82 | RM4-5 | Invitrogen |
| NKp46 | AF2225 | | R&D |
| HMGB1 | 651406 | 3E8 | Biolegend |
| Goat IgG | A11055 | | Invitrogen |
| Rabbit IgG | A31573 | | Invitrogen |
| Hoecht | 40046 | | Biotium |

### Supplementary Table 4. The list of RT-qPCR primers

| **Target gene** | **Forward Primer (5’ to 3’)** | **Reverse Primer (3’ to 5’)** |
|---|---|---|
| mSlurp1 | GGTCACGGAAGCAACAGAAG | GGCCTTCCGATGCTATACCT |
| mKlf4 | TGCCAGACCAGATGCAGTCAC | GTAGTGCCTGGTCAGTTCATC |
| mCd200 | ACAGCCCATAGTACACCTTCA | TGTCCCAGTACCCTTCCAGG |
| mIfngr1 | TACAGGTAAAGGTGTATTCGGGT | ACCGTGCATAGTCAGATTCTTTT |
| mGapdh | AATGTGTCCGTCGTGGATCTGA | GATGCCTGCTTCACCACCTTCT |

### Supplementary Table 5. The list of shRNA sequences

| shRNA | Catalog # | shRNA sequence |
|---|---|---|
| shScrambled | TRCN0000190210 | shRNA Control Plasmid |
| shSlurp1 #1 | TRCN0000190210 | CCTGTAAGACTGTACTGGAGA |
| shSlurp1 #2 | TRCN0000190254 | CTTCCGATGCTATACCTGTGA |
| shSlurp1 #3 | TRCN0000189776 | GAAGACACAGCCTGTAAGACT |
| shIfngr1 #1 | TRCN0000067368 | CCACATAGAATATCAGACTTA |
| shIfngr1 #2 | TRCN0000067369 | GCCAGAGTTAAAGCTAAGGTT |
| shIfngr1 #3 | TRCN0000067371 | CCACACTGGATTCCAGATATT |
| shCd200 #1 | TRCN0000066679 | GCCCATAGTACACCTTCACTA |
| shCd200 #2 | TRCN0000066681 | CGAGAGTCACTTCCATTCAAA |
| shCd200 #3 | TRCN0000066682 | CAGAGTCTGGACAAAGGATTT |
| shCd200 #4 | TRCN0000066678 | CCTGCCTACAAAGACAGGATA |
| shKlf4 #1 | TRCN0000095370 | CTCTCTCACATGAAGCGACTT |
| shKlf4 #2 | TRCN0000095371 | CTGGACCTAGACTTTATCCTT |

Supplementary Table 6. The list of ChIP qPCR Primers

| Slurp1 promoter region | Forward Primer (5' to 3') | Reverse Primer (3' to 5') |
|---|---|---|
| -850 ~ -600 | CCATCCACAGGGCCACTCAT | GTCCTGACAGCAGAACTCTACCT |
| -500 ~ -250 | CAGGTACTCCCTCCTTTCCATACTG | ACTGAGGAAGCCTTTTAGAGCC |
| -150 ~ +30 | GGCCCCACCCTGGGATGGTAGGTGA | TCTTCAGTGCTCAGGAGCTAGGA |

| Cd200 promoter region | Forward Primer (5' to 3') | Reverse Primer (3' to 5') |
|---|---|---|
| -200 - +30 | CTACGTCACCCTATACTGCCATTTGG | AGGCAGACTCTACAGCTCCTCTAGTG |
| -500 - -300 | CAGGTACTCCCTCCTTTCCATACTG | ACTGAGGAAGCCTTTTAGAGCC |
| -800 to -600 | GGCCCCACCCTGGGATGGTAGGTGA | TCTTCAGTGCTCAGGAGCTAGGA |
| -1100 to -900 | GAATACCTTCTCACACCAGAGAGACTAG | TCGAGAAAGAGAGAGAGAGGAGAGAG |

Supplementary Table 7. Slurp1 sgRNA target sequences and PCR primer sequences used in Slurp1 KO validation

| **Name** | **Forward sequence (5' to 3')** | **Reverse Sequence (3' to 5')** |
|---|---|---|
| Slurp1 sgRNA #1 | CCATCCACAGGGCCACTCAT | GTCCTGACAGCAGAACTCTACCT |
| Slurp1 PCR #1 | GCCAGGCTCTAAAAGGCTT | CTGTCTCCAGTACAGTCTTAC |
| Slurp1 PCR #2 | GACAGCAGAGCATGGTGTC | GTCTTCCATCTTGCACTGAG |
